# Single genome analysis of normal cells identifies the early steps of kidney cancer

**DOI:** 10.64898/2026.09.03.749153

**Authors:** Jean Piero Margaria, Giovanni Battista Pipitone, Alessandro Larcher, Sipontina Faienza, Isaline Rowe, Francesca Corea, Valentina Piatti, Nicola Sala, Jose Manuel Garcia Manteiga, Jianfeng Ge, David Mas-Ponte, Gianfranco Distefano, Sakari Vanharanta, Umberto Capitanio, Fran Supek, Paola Carrera, Andrea Salonia, Alessandra Boletta, Irene Franco

**Affiliations:** Somatic Mutation Mechanisms Unit, Division of Genetics and Cell Biology, IRCCS Ospedale San Raffaele, Italy; Vita-Salute San Raffaele University, Milan, Italy; Genomics for Diagnosis of Human Disease Unit, IRCCS Ospedale San Raffaele, Milan, Italy; Medical Genetics Unit-Molecular Genetics Laboratory, IRCCS Ospedale San Raffaele, Milan, Italy; Division of Experimental Oncology/Unit of Urology; URI; IRCCS Ospedale San Raffaele, Milan, Italy; Center for Omics Sciences, IRCCS Ospedale San Raffaele, Milan, Italy; Early Cancer Institute, University of Cambridge, Cambridge, UK; Institute for Research in Biomedicine (IRB), Barcelona, Spain; Cystic Kidney Disorders Unit, Division of Genetics and Cell Biology, IRCCS Ospedale San Raffaele, Milan, Italy; Translational Cancer Medicine Program, University of Helsinki, Helsinki, Finland; Biotech Research and Innovation Centre (BRIC), University of Copenhagen, Copenhagen, Denmark

## Abstract

Whole genome analyses have shown that every cell of the body accumulates thousands of genetic changes during a lifetime, with implications for cancer initiation. We have analyzed normal kidney cells from both healthy individuals and patients with the Von Hippel Lindau (VHL) syndrome, which predisposes to kidney cancer. Using a clonal-expansion method that enables the collection of high confidence genetic and gene expression data from the same cell, we define a new mutational signature that marks a specific subset of damage-activated proximal tubule cells, as well as 96% of clear cell renal cell carcinomas. The underlying mutational process is specifically active in highly transcribed DNA and impacts the occurrence of cancer driver variants in the *VHL* gene. Our data from normal cells, supported by *in vitro* experiments, point to perturbations of oxygen availability as a causative factor. These findings link hypoxia, a frequent endogenous stress in the normal renal tissue, to excessive mutation and cancer initiation.

## Introduction

The complex morphology of the kidney is designed to maintain the body balance of fluids and electrolytes. The kidney tubule is functionally subdivided in segments, each one specialized in the exchange of water and solutes with the pre-urine. A specific segment, namely the proximal tubule (PT), is the most common site for tumor formation ^1–5^. The factors that favor cancer incidence in the PT are still not known, but an emerging hypothesis is that the specific activities of the PT may expose it to tissue and DNA damage, which contribute to the tumorigenic process ^3,6–9^.

Single cell RNAseq studies have supported the annotation of more than 20 types of fully differentiated, post-mitotic cells in the renal tubules. Moreover, a rich landscape of cellular states representing damaged, damage-activated, and proliferating kidney epithelial cells has been described ^10–12^. Functional studies have shown that tubular damage activates proliferation in surrounding cells by inducing both the expansion of resident progenitor cells and the de-differentiation of fully differentiated tubular cells ^13–15^. Activated cells are rare in the human kidney, but consistently found in healthy individuals, suggesting that the tissue undergoes damage-repair cycles, even in individuals without clinical symptoms ^11,12^. In the urine, the concentration of proliferation-competent cells is higher compared to kidney biopsies, and drastically increases upon an acute kidney injury episode ^16–18^. Cells in the activated state upregulate proliferative and embryonic programs ^10^, typically associated with tumorigenesis ^3^. In agreement, a rare PT-resident cell subset expressing the embryonic nephrogenic factor *PAX2* and the inflammatory marker *VCAM1* has been proposed as the cell of origin of clear cell (ccRCC) and papillary (pRCC) renal cell carcinoma ^5,19^.

The acquisition of cancer-driver genetic variants is part of the process of transformation of a normal cell into cancer and is commonly observed in ccRCC ^20^. The Von Hippel Lindau (*VHL)* tumor suppressor is the most commonly mutated gene (>80% of ccRCC cases) ^20,21^ and *VHL* loss is recognized as the key genetic alteration that initiates tumorigenesis in this tumor subtype ^22^. In the inherited VHL syndrome, caused by germline, loss-of-function mutations in one allele of *VHL*, patients develop multiple ccRCCs, with 70% penetrance and early onset (mean age of onset 44 *vs* 65 in the general population) ^23^.

Therefore, we sought to analyze the genome of proliferating kidney tubule cells from both healthy and VHL-disease individuals and to investigate whether the transition from normal to cancer is accompanied by specific mutational processes. In particular, we analyzed thousands of single base substitutions and insertion-deletions detected at whole genome level in kidney cells to identify specific mutational signatures. The latter are signals of active mutational processes in the cell lineage history and have been widely analysed in cancer samples as a source of information about the life-long exposure to mutagens ^24–26^. A recent analysis of more than 900 ccRCC genomes has allowed a deeper characterization of the mutational signatures active in PT-derived kidney cancers and their cell of origin ^27^. This study has identified a highly prevalent single base substitution signature named SBS40b, which a) is restricted to cancers derived from the PT; b) presents a geographic distribution that correlates with kidney cancer risk; c) correlates with blood indicators of kidney damage ^27^, overall highlighting a potential causal link between PT damage and cancer-causing mutation in the kidney. This link is strengthened by an analysis of normal kidney tubule cells, which has identified a population of damage-activated PT cells that showed an excessive mutation load, mostly due to the activity of the SBS40 signature ^28^.

Following up on this study, we have established a protocol that enables the collection of high confidence somatic variants and gene expression data from the same normal kidney cell. The method, based on *in vitro* clonal expansion, restricts the analysis to proliferation-competent cells. Importantly, we applied this protocol to non-invasive urine samples, and demonstrated that we can retrieve the same populations of proliferating cells from both kidney biopsies and urine samples. Cell collection from the urine enables the analysis of a wide range of individuals, including healthy volunteers and individuals with the VHL syndrome. By analyzing normal cells and renal cancers, we have found a specific mutational process that increases mutation burdens, particularly in highly transcribed DNA, and favors the occurrence of cancer driver variants. Our study shows that this cancer-causing mutational process is induced by environmental factors. Nonetheless, complete loss of pVHL also compromises genome integrity. Overall, our study sheds light on cancer initiation in human kidney tissue with a mechanistic link.

## Results

### Concomitant gene-expression and somatic mutation profiling of clonally expanded, normal kidney cells

To characterize the age-related somatic mutation in human kidney cells, we developed a protocol that enabled the concomitant collection of somatic mutation and gene expression data from the same cell (Fig 1a). We used an *in vitro* clonal expansion strategy, a golden standard method for acquiring high genome coverage and high confidence somatic mutation data from single somatic genomes ^29^. We obtained kidney cells from urine samples of 9 healthy volunteers (*VHL+/+*, age range 24-70) and 9 individuals with the VHL hereditary cancer predisposition syndrome (*VHL+/−,* age range 29-56, Suppl Table 1 and 2). Moreover, we obtained cells from biopsies of normal kidney parenchyma collected from patients diagnosed with ccRCC and elected for surgical treatment with nephrectomy (1 *VHL+/+* and 2 *VHL+/−* kidneys explanted for oncological treatment, Suppl table 1 and 2).

**Figure 1.**
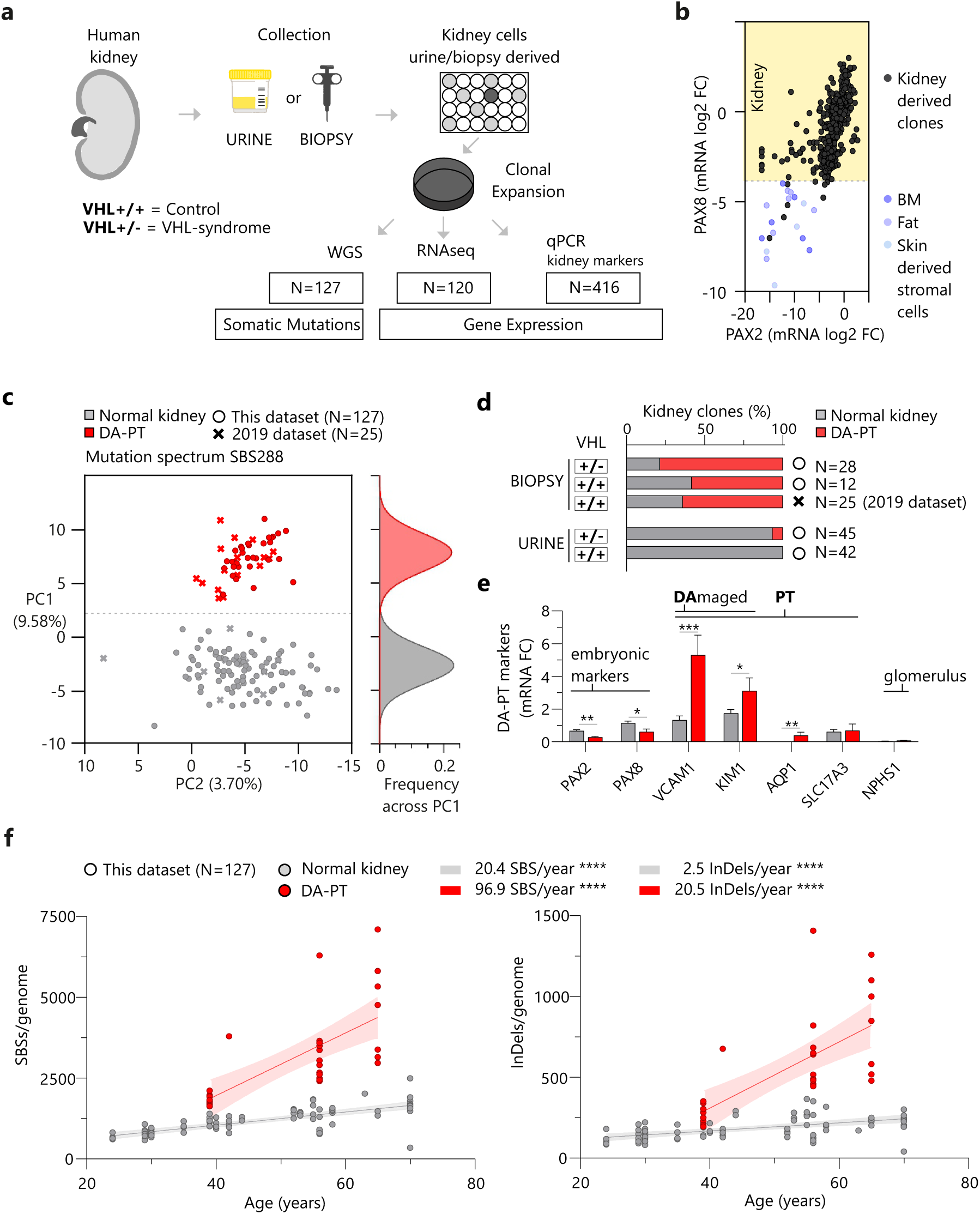
Single-genome analysis of normal kidney cells from controls and patients with inherited VHL (Von Hippel Lindau) disease. **a.** Schematic representation of the method used for concomitant analysis of gene-expression and somatic mutations in clonally expanded, human kidney cells of either control individuals (*VHL+/+,* n=10) or VHL-syndrome patients (*VHL+/−,* n=9). Kidney cells were grown from either kidney biopsies or urine samples. All clones (n=416) were tested for the expression of renal markers using qPCR, and 120 clones were also tested by RNAseq. N=127 clones were subjected to whole genome sequencing (WGS), to detect high confidence somatic variants accumulated *in vivo* in the single cell that initiated the colony. Matching WGS, RNAseq and qPCR data were available for n=56 clones. **b.** Expression of the nephrogenic lineage markers *PAX8* and *PAX2* in single clones (n=416), compared to the expression levels quantified in cultured stromal cells derived from extrarenal tissues (bone-marrow derived mesenchymal cells, skin fibroblasts, fat progenitors). Expression levels were quantified by qPCR and reported as relative to a standard sample (normal kidney tissue biopsy). **c.** Principal Component Analysis of SBS288 mutation spectra in single, kidney-derived normal genomes sequenced in either this study (n=127) or a previous study (2019 dataset, n=25). The distribution curve of two identifiable clone clusters along the PC1 is shown on the right. **d.** Percentage of *VHL+/+* and *VHL+/−* clones belonging to the two clusters defined in Figure 1c (*damaged activated (DA)-PT* and *normal kidney cells*) observed in either urine or kidney samples. **e.** Average expression levels of renal markers in the the *DA-PT* (n=32) versus all remaining *normal kidney clones* (n=120). **f.** Number of somatic single base substitutions (SBSs)(left panel) and insertions/deletions (InDels)(right panel) per genome and linear regression with age of the donor in *DA-PT* and *normal kidney clones*. * p<0.05; ** p<0.01; *** p<0.005, by Student’s t test or simple linear regression to measures a significantly non-zero slope as appropriate.

Freshly isolated single cells were clonally expanded and subjected to qPCR analysis of kidney marker genes, including *PAX8* and *PAX2,* two embryonic, nephric-lineage specific, transcription factors (Suppl Data 1). In mature adult tissue, *PAX8* is specifically expressed by kidney epithelial cells of all segments of the nephron, but absent in stromal and endothelial cells ^12,30^. Instead, *PAX2* is silent, but re-activated by proliferating cells scattered along the nephron ^12,31,32^. A total of 416 clones were tested. In 98.6% of clones, the levels of *PAX8* were higher than those measured in stromal cells from non-kidney tissues, while 91.8% of clones expressed higher levels of both *PAX2* and *PAX8* (Fig 1b), overall validating our clones’ origin from proliferating kidney epithelial cells. In further support, RNAseq data of selected clones (n=120) were compared with published scRNAseq datasets from human kidneys and urine^12,17^ (Suppl Data 2). Clones were largely positive for transcriptional signatures of various subsets of proliferating cells of the kidney tubule, including the VCAM1-PT (proximal tubule) cells from the kidney ^12^ and the inflamed-, damaged- and proliferative-tubule epithelial cells (TEC) from the urine ^17^. In contrast, the signatures of fully differentiated cells were mostly negative (Suppl Fig 1a-b). When we compared the populations of clones derived from either urine *vs* kidney biopsies, or *VHL+/−* vs *VHL+/+* individuals, we found a broad overlap (Suppl Fig 2). Differences in specific genes tested by qPCR are shown in Suppl Fig 3.

Next, we did whole genome sequencing (WGS) of clone-derived DNA and obtained single genome, somatic mutation data for n=54 *VHL+/+* and n=73 *VHL+/−* clones (Suppl Data 3). We integrated the dataset with 25 *VHL+/+* clones from our published normal kidney clone study (identified as “2019 dataset”) ^28^ and performed mutation spectra analyses on a total of n=152 genomes (Fig 1c). First, we applied a classification of single base substitutions (SBSs) that takes into account the type of base substitution, the trinucleotide context, the position of each mutation in genic or intergenic DNA, and the direction of transcription, ultimately recognizing 288 types of SBSs (SBS288). Principal component analysis (PCA) of the SBS288 spectra highlighted a specific cluster of genomes that showed high principal component 1 (red cluster, Fig 1c). These clones were the most frequent type among biopsy-derived clones, but they were rare in urine (Fig 1d). We named this cluster *damage-activated PT (DA-PT)*, as it was characterized by higher expression of proximal tubule and PT-damage markers (*VCAM1*, *KIM1, AQP1)*, compared to the remaining clones, identified as *normal kidney* (Fig 1e, qPCR quantification of specific genes; Suppl Fig 4a-b, transcriptional signature scores from RNAseq data).

DA-PT genomes presented high-for-age mutational burdens of both SBSs and small insertions-deletions (indels; Fig 1f). We performed a mutational signature analysis and fitting to COSMIC signatures with SigProfiler ^33^. This analysis included somatic mutation data from extra-renal samples available in the literature ^28,34,35^ and 10 ccRCC genomes from 3 VHL-disease patients included in the study (Suppl Data 4). The main signatures in kidney-derived clones were COSMIC SBS signatures 1, 5, 8, 40b and 40c. The SBS40b was the distinctive signature found in ccRCC, with 8/10 positive genomes. Interestingly SBS40b was also found in 24/48 DA-PT genomes and none of the normal kidney genomes (Suppl Fig 5a). Signature extraction with a different method confirmed the SBS40b signal in all samples identified by SigProfiler (Suppl Fig 5b). In addition, DA-PT cells showed an enrichment of 1 bp T deletions and 1 bp T insertions at 2-to-5 bp-long T homopolymers, compared to the normal kidney genomes, and a similar indel pattern was observed in ccRCC (Suppl Fig 6a-b). Despite these similarities with ccRCC, DA-PT genomes showed lower mutational burdens (Suppl Fig 6c) and lacked the typical ccRCC driver mutations, i.e. chromosome 3p loss, second-hit mutations in *VHL*, and extensive copy number alterations, that were observed in our ccRCCs genomes (Suppl Fig 7). Overall, our analysis supports the idea that DA-PT clones are not derived from cancers, but are a subset of normal cells exposed to a specific mutational process.

Finally, we assessed the effect of the germline loss-of-function mutation in one allele of *VHL* by comparing the somatic variants found in *VHL+/+* and *VHL+/−* individuals (Suppl Fig 8). No differences were noticed when plotting the PC1 and PC2 of the SBS288 mutation spectrum (Suppl Fig 8a). However, the *VHL+/−* DA-PT clones showed higher incidence of COSMIC SBS40b, and lower incidence of SBS40c, compared to their *VHL+/+* counterpart (Suppl Fig 8b). When we assessed the mutational burdens (Suppl Fig 8c-d), we did not find significant differences, but an interesting trend towards an increased yearly rate of indel accumulation in *VHL+/− normal kidney* cells (Suppl Fig 8d). We could not assess the difference in *DA-PT*, as they were derived from two different datasets, characterized by slightly different mutation counts.

In summary, our analysis identifies a subset of normal kidney cells that present an excess of somatic variants and a specific SBS spectrum, together with high levels of transcriptional markers of PT-damage. These cells can be grown from kidney biopsies, as well as from urine samples.

### *DA-PT* cells are subjected to a transcription-coupled mutational process (ToCCATA)

We leveraged our somatic mutation data to obtain insight into possibly dysregulated DNA damage/repair mechanisms in the *DA-PT cells*. A distinctive feature of somatic mutation in *DA-PT* cells was an excess of mutations in genic regions (Fig 2a). Notably, mutations increased along with gene expression levels in *DA-PT cells* (Fig 2b), while *normal kidney cells* displayed the opposite trend, with highly expressed DNA protected from mutation (Fig 2b). These patterns highlight an opposite effect of transcription on DNA damage and repair in *DA-PT* and *normal kidney cells*.

**Figure 2.**
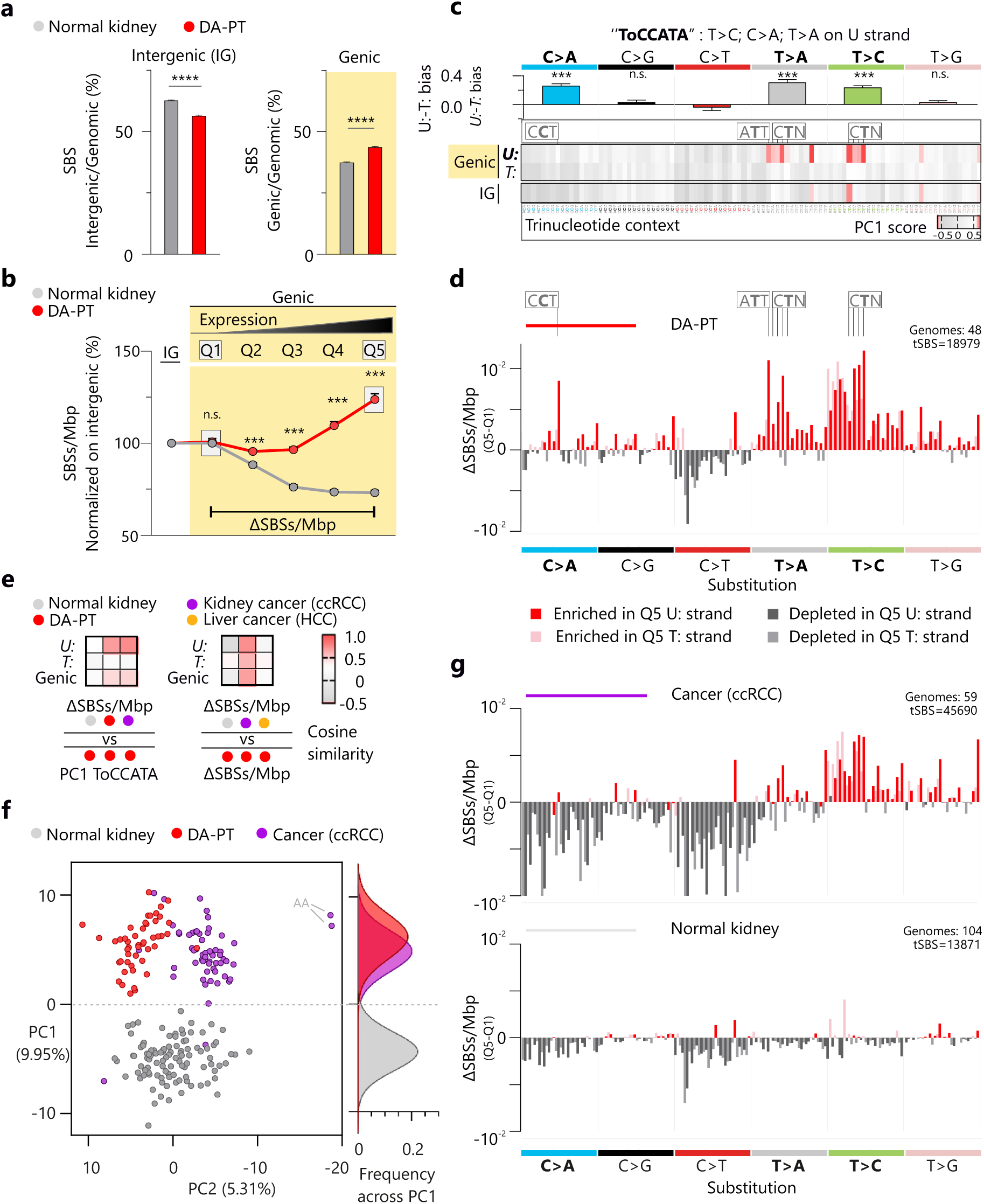
A transcription-coupled mutational process induces the ToCCATA (T>C;C>A;T>A variants on U strand of genic DNA) mutational signature specifically in DA-PT and ccRCC genomes. **a.** Distribution of SBSs in either intergenic or genic DNA in *DA-PT* and *normal kidney* clones. **b.** Density (mutations/Mbp) of SBSs in either intergenic or genic DNA, subdivided in 5 bins according to gene expression levels obtained from a curated list (Human protein atlas) of genes expressed in healthy human kidney. **c.** Heatmap depicting the influence of every SBS288 class on the principal component 1 (PC1) of the PCA presented in Fig 1c. The reported U: to T: bias is the mean and s.e.m. of 16 trinucleotide-context specific U: to T: ratios per substitution type. **d.** Spectrum of SBSs enriched in highly expressed (Q5) compared to non-expressed genic DNA (Q1 of Fig 2b) in aggregated somatic mutation data from *DA-PT* genomes. **e.** Cosine similarity scores from the comparison of the SBS spectra in Fig 2c, 2d and 2g. **f.** Principal component analysis of SBS288 mutation spectra in single, *normal kidney* and *DA-PT* genomes, together with kidney cancer (ccRCC) genomes from either VHL-disease patients (n=10) or the PCAWG dataset (N=49). Two kidney cancer samples showing exposure to aristolochic acid, a known renal mutagen, are indicated with the acronym “AA”. **g.** Spectrum of SBSs enriched in highly expressed (Q5) compared to non-expressed genic DNA (Q1) in aggregated somatic mutation data from kidney cancer (ccRCC) and normal kidney genomes. *** p<0.005, by Student’s t test or 1 or 2-way ANOVA followed by Holm-Šídák’s multiple comparisons test as appropriate.

We sought to understand what components of the mutation spectrum were induced by the transcription-coupled mutation process. We applied two different analytical methods, which provided overlapping results (Fig 2c-e). First, we plotted the components of the SBS288 spectrum that guided the separation of the *DA-PT* from the *normal kidney* genomes along the principal component 1 of Fig 1c (methods). The SBS types that characterized the *DA-PT* clones were mostly in genic DNA, particularly in the un-transcribed strand (Fig 2c, red shades of the heatmap). We named this pattern ToCCATA, from its distinctive feature of transcriptional strand bias in T>C, C>A, T>A substitutions (Fig 2c, top). The spectrum did not coincide with the spectrum of SBS40b (Suppl Fig 9a). However, ToCCATA and SBS40b exposure showed a significantly positive correlation in our dataset (Suppl Fig 9b-c), suggesting that these two mutational patterns might share a common etiology.

The second method to derive the components of the transcription-induced mutation spectrum observed in *DA-PT cells* identified the SBS types that were more abundant in the highly (Q5) versus lowly expressed (Q1) portions of the transcribed genome in *DA-PT* cells in Fig 2b (ΔSBS/Mbp, see methods). The profile of U: strand SBSs enriched in the top expressed quintile of *DA-PT* cells (Fig 2d) highly resembled the spectrum of ToCCATA identified by PCA in Fig 2c (cosine similarity 0.78, Fig 2e). In contrast, the profile of T: strand SBSs was mildly correlated (cosine similarity 0.29, Fig 2e). Interestingly, the spectrum had a clear dependency on gene expression levels (Suppl Fig 10a-c). Moreover, the spectrum observed in *DA-PT cells* was different from the spectrum of another transcription-coupled mutational process, previously reported in hepatocellular carcinoma ^36^ (Suppl Fig 10d, cosine similarity −0.27, Fig 2e). Overall, these results prove that ToCCATA is a newly reported mutational signature, induced during transcription.

Here we report that *DA-PT* cells are subjected to a transcription-coupled mutational process which induces T>C, C>A, T>A substitution classes, specifically in the untranscribed DNA strand (ToCCATA).

### ToCCATA-bearing, DA-PT cells are the origin of ccRCC

At this point, we sought to analyze the mutation spectrum of PT-derived cancers. We derived the SBS288 spectra of 10 ccRCCs from our VHL-disease patients (hereditary ccRCC), together with an additional n=49 sporadic ccRCCs from PCAWG ^37^. Again, *normal* and DA-PT *kidney* genomes separated along the principal component 1 axis. Interestingly, the majority of cancers (96%) clustered together with *DA-PT kidney* genomes (Fig 2f, Suppl Fig 11a). In line with this finding, the profile of SBSs enriched in the top expressed quintile (Q5 – Q1) of ccRCCs was similar to the profile observed in *DA-PT* (Fig 2e and g), while normal kidney cells showed a depletion of most mutation types in highly expressed genes (Fig 2e and g). This pattern did not depend on *VHL* status (Suppl Fig 11b). Overall, these results indicate that *DA-PT* genomes and ccRCC samples share a history of exposure to the same mutational process, which produces the ToCCATA mutational pattern.

In further proof, we assessed the ToCCATA index, which summarizes the ToCCATA spectrum to a single value (the difference between U: and T: T>C, C>A, T>A variants, normalized on the total number of variants) and offers the advantage of measuring ToCCATA in single, rather than in aggregated genomes. The ToCCATA index correctly discriminated *normal kidney* cells (negative values) from *DA-PT* cells (positive values) (Fig 3a, Suppl Data 3). CcRCC genomes of our VHL-disease patients also showed high ToCCATA index, and this trend was even more pronounced when the analysis was restricted to likely clonal mutations (Fig 3a, see methods). The latter observation supports that ToCCATA occurs before cancer transformation.

**Figure 3.**
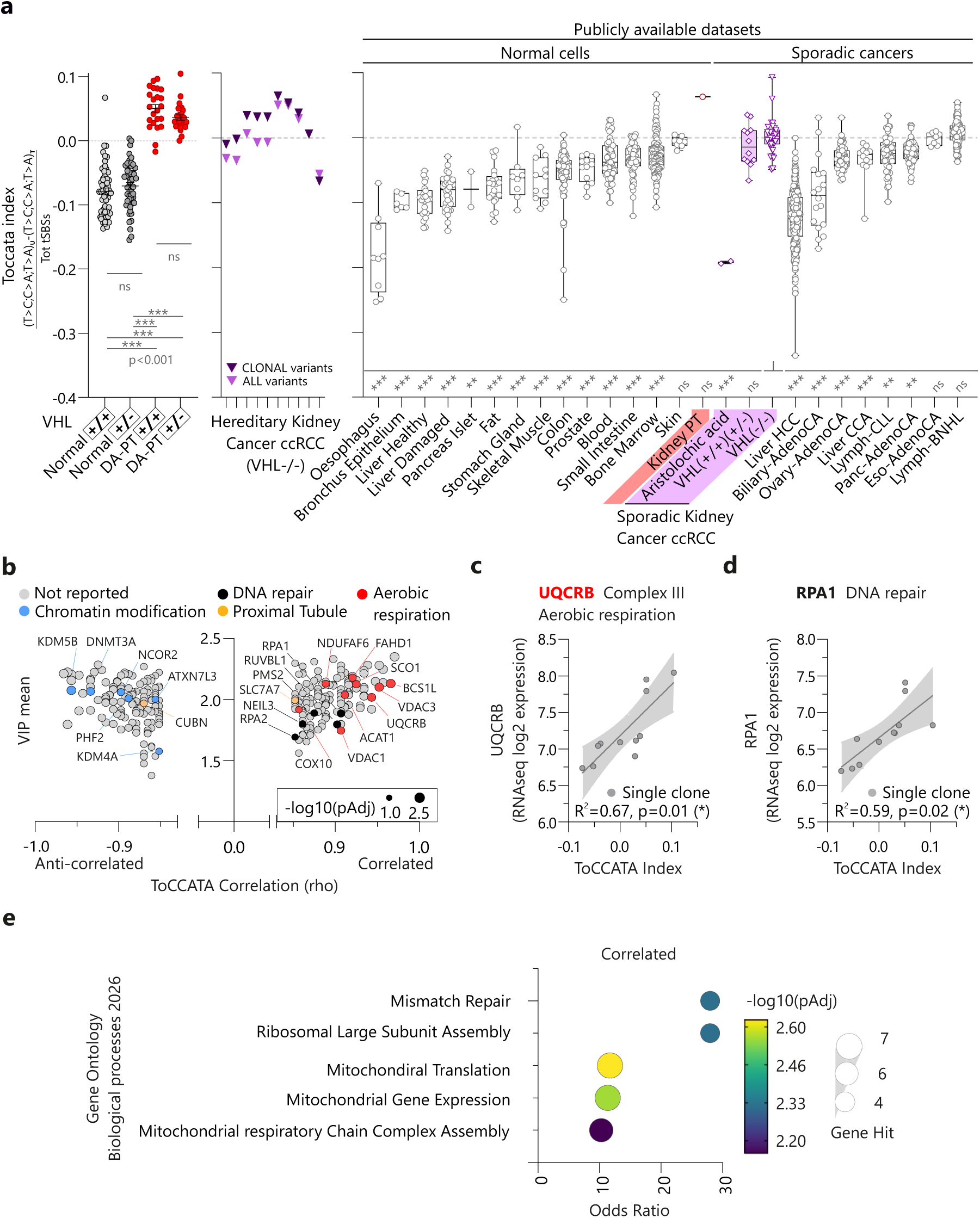
ToCCATA correlates with enhanced mitochondrial metabolism. **a.** ToCCATA index (i.e. transcriptional strand bias of selected substitution types: T>C; C>A; T>A) in either kidney genomes from Fig 1c (left panels) or a collection of published single genome datasets from normal tissues and sporadic cancers (right panel, listed in Suppl Data 5). Single sporadic kidney cancer (ccRCC) genomes were divided into 3 groups: exposed to aristolochic acid (AA), intact *VHL* (*VHL+/+; VHL+/−)*, biallelic loss of *VHL* (*VHL−/−*). Publicly available datasets were all compared with sporadic kidney cancer VHL−/− group. ** p<0.01; *** p<0.005, by 1-way ANOVA followed by Holm-Šídák’s multiple comparisons test. **b.-e.** Results of the Spearman correlation of the ToCCATA index and the expression of single genes from RNAseq data in 11 clones from a single individual (*VHL+/−,* aged 56). Volcano plot of the correlated and anti-correlated genes with significant BH adjusted p value **(b).** Linear correlation of two representative genes with ToCCATA. * p<0.05; by simple linear regression; we tested the significant deviation of the slope from zero. (**c-d**). Biological processes that showed a BH adj p <0.01 enrichment, using the Gene Ontology database **(e).**

Next, we investigated the tissue distribution of ToCCATA using published datasets. We compiled a somatic mutation dataset comprising single genomes derived from several types of normal tissues and sporadic tumors that presented >1000 SBSs/genome (Suppl Data 5). The ToCCATA index discriminated the only available PT cell genome from the other normal genomes, suggesting that ToCCATA index >0 is a feature specific to the kidney tissue and restricted to the *DA-PT* subset. In addition, a positive ToCCATA index was found in most ccRCCs from the PCAWG dataset, except for those tumors that bore the signature of aristolochic acid (AA), a known mutagen that induces T>A mutations biased towards the transcribed strand, i.e., opposite to that observed in *DA-PT* cells.

In summary, we did an analysis of mutational patterns, focusing on the effect of transcription. We found a transcription-coupled mutational process that specifically affects the *DA-PT* cells. This process induces a mutational signature that we named “ToCCATA”. Remarkably, ToCCATA is found in 96% of ccRCC genomes analyzed, more prominently in clonal variants, overall suggesting that ccRCC originates from the subset of cells that we have defined as *DA-PT* cells.

### In vitro modeling of PT-damage by exposure to hypoxia promotes ToCCATA

To understand what process might be the trigger of ToCCATA, we leveraged our coupled somatic mutation and gene-expression data from single clones. We scored the genes whose expression significantly correlated with the ToCCATA index in our dataset (n=56 clones, Suppl Data 6). In support of the PT-identity of ToCCATA-bearing cells, the expression of the PT-specific transporters SLC2A9 and SLC7A7 positively correlated with ToCCATA (Adj p value 0.0015 and 0.0004, respectively; Suppl Data 6). Moreover, in line with the high mitochondrial activity of differentiated PT cells, the genes that positively correlated with ToCCATA were significantly enriched for the mitochondrial compartment (SubCell BarCode, Odds ratio 9.73, Adj p value 0.00008). These data were largely replicated in intra-individual correlations (VUD12=15 clones; VUD1=11 clones, Suppl Data 6). For individual VUD1, the correlation was particularly informative (Fig 3b-e). The genes that positively correlated with ToCCATA were enriched for mitochondrial processes, including aerobic respiration (Fig 3b-c-e). ToCCATA-positive cells showed an upregulation of DNA repair components (Fig 3b-d-e). In contrast, multiple chromatin modifiers negatively correlated with ToCCATA, including the oxygen-dependent histone demethylases KDM5B and KDM4A (Fig 3b).

Since oxygen metabolism is a key feature of ToCCATA bearing cells, we hypothesized that perturbations of mitochondrial respiration could alter cellular homeostasis and culminate in excessive mutation. Indeed, the PT is one of the most oxygen-demanding tissues in the body and is very sensitive to ischemic insults ^38^. In addition, the *VHL* gene, which is commonly mutated in PT-derived tumors, encodes for a negative regulator of metabolic adaptation to low oxygen ^20^. Therefore, we tested whether a sudden perturbation of mitochondrial metabolism, due to either exposure to insufficient environmental oxygen (hypoxia) or genetic loss of *VHL* (pseudohypoxia, schematics in Fig 4a) could stimulate ToCCATA-related mutagenesis *in vitro*.

**Figure 4.**
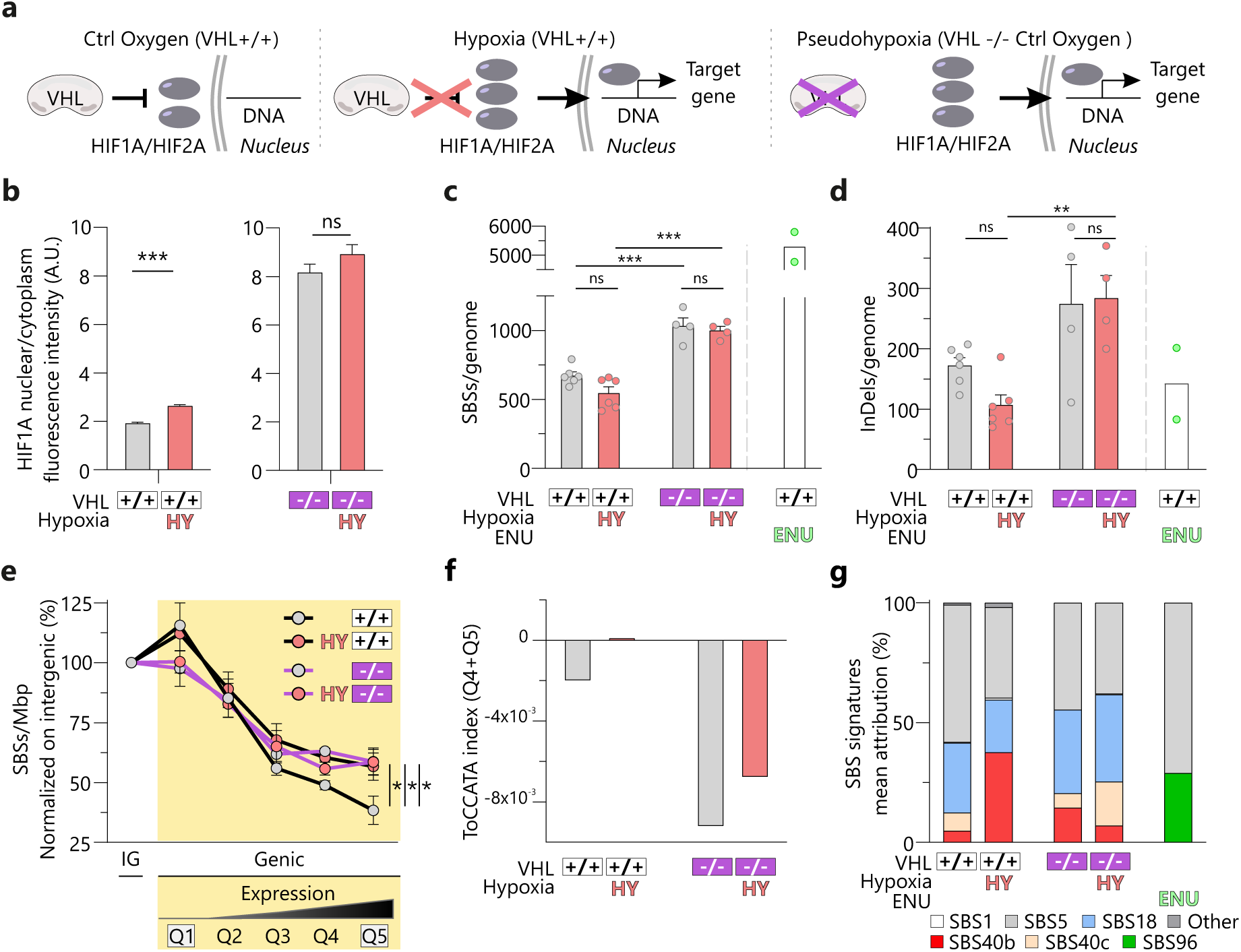
Environmental hypoxia induces ToCCATA/SBS40b in a PT cell line. **a.** Schematic representation of the VHL/HIF pathway in normal (**left**) and low oxygen conditions (hypoxia, **middle**), and similar signaling upon genetic loss of *VHL* (pseudohypoxia, **right panel**). **b.** Quantification of HIF1A nuclear/cytoplasmic immunofluorescent signal in engineered HK2 cells upon hypoxia (**left**) and genetic loss of *VHL* (**right**). **c**.-**g**. Analysis of *in vitro* mutations detected in single genomes and grouped by treatment (VHL-WT ctrl oxygen n=6; VHL-WT hypoxia, n=6; VHL-KO ctrl oxygen, n=4; VHL-KO hypoxia, n=4 single genomes). Total number of SBSs (**c**) and indels (**d**) per genome; density (mutations/Mbp) of SBSs in either intergenic or genic DNA, subdivided in 5 bins according to gene expression levels obtained from RNAseq data of VHL-WT and VHL-KO ctrl oxygen HK2 cells. *p<0.05 by two-way ANOVA followed by Dunnett’s multiple comparisons test against the control oxygen VHL+/+ condition (**e**); ToCCATA index (transcriptional strand bias of T>C; C>A and T>A) of the mutations found in highly expressed DNA (Q4 and Q5) (**f**); COSMIC signature relative exposure (**g**). Two VHL-WT clones exposed to ENU treatment in control oxygen are shown as a reference and not included in the statistical analyses. **p<0.01, ***p<0.005 by Student’s t-test or one-way ANOVA followed by Holm-Šídák’s multiple comparisons test, as appropriate.

We used an immortalized cell line derived from normal human proximal tubule (HK2) and engineered to conditionally express VHL ^39^, and performed the experiments in the acute phase after cellular depletion of the VHL protein (Suppl Fig 12a). We exposed VHL-WT and VHL-KO cells to either a control (22% oxygen) or hypoxic (0.1% oxygen) cell culture for 72 hours. We verified the expected activation of the HIF pathway in low oxygen by measuring HIF1a levels and translocation to the nucleus (Fig 4b, Suppl Fig 12b-d). Subsequently, we identified the *in vitro* accumulated SBSs and indels in individual cells, upon re-cloning and WGS (n=4-6 clones per condition). The VHL-KO cells displayed a general increase of *in vitro* acquired SBSs and indels compared to their VHL-WT counterpart (Fig 4c-d), indicating that pVHL is needed for protection from mutation in cultured cells. The loss of *VHL* altered the mutational patterns, but there was no additional alteration induced by exposure to hypoxia in VHL-KO cells (Fig 4e-g, Suppl Fig 12e), in line with the pattern of HIF activation (Fig 4b). Conversely, when VHL-WT cells were exposed to hypoxia, *in vitro* mutations accumulated in highly-expressed genes (Fig 4e) and the transcriptional strand bias was in favor of the un-transcribed strand (positive ToCCATA index, Fig 4f), two distinctive features of ToCCATA. When we analyzed COSMIC SBS signatures, we found that environmental hypoxia increased the incidence of SBS40b (Fig 4g, Suppl Fig 12f). VHL-loss did not have a clear effect on SBS40b, but increased the incidence of SBS18 (induced by supra physiological oxygen levels in control cell culture and characterized by C>A variants in T strand, Suppl Fig 12g), possibly affecting the quantification of the ToCCATA index (Fig 4e). Overall, this experiment points to adaptation to hypoxia as a stimulus able to induce mutations of the ToCCATA spectrum.

In summary, we show that both genetic loss of *VHL* and environmental hypoxia promote mutation *in vitr*o, but the ToCCATA mutational pattern is specifically stimulated by environmental hypoxia in *VHL*-competent cells. These data suggest that the response to environmental hypoxia could be the cause of accelerated mutation accumulation in highly expressed genes observed *in vivo* in *DA-PT* cells.

### The ToCCATA signature influences the appearance of cancer driver variants in the VHL gene

Our data show that the ToCCATA signature is present in kidney pre-cancer cells and exacerbated in highly expressed genic regions (Fig 2), thus we asked whether ToCCATA functionally contributes to cancer initiation. The *VHL* gene, harboring cancer driver variants in more than 80% of sporadic kidney cancers ^21^, is robustly expressed in normal kidney. To evaluate the contribution of ToCCATA to the occurrence of *VHL* kidney cancer driver variants, we collected all missense variants that have been reported as cancer drivers in the *VHL* gene in the AACR Project GENIE consortium database ^40^. We plotted the SBS288 spectra for these variants, dividing those found in ccRCC (exposed to the mutational process underlying ToCCATA) from those found in non-kidney cancer types (not exposed, Suppl Fig 13a). We noticed that ccRCCs showed a significantly higher incidence of ToCCATA-related mutation classes (T>C, C>A, T>A substitutions on U strand) (Fig 5a). Similarly, we highlighted a positive correlation between the spectrum of *VHL* cancer-driver variants and the spectrum of ToCCATA, exclusively for ccRCC (Fig 5b), supporting the idea that exposure to the mutational process underlying ToCCATA significantly shapes the spectrum of cancer driver variants observed in ccRCC.

**Fig. 5:**
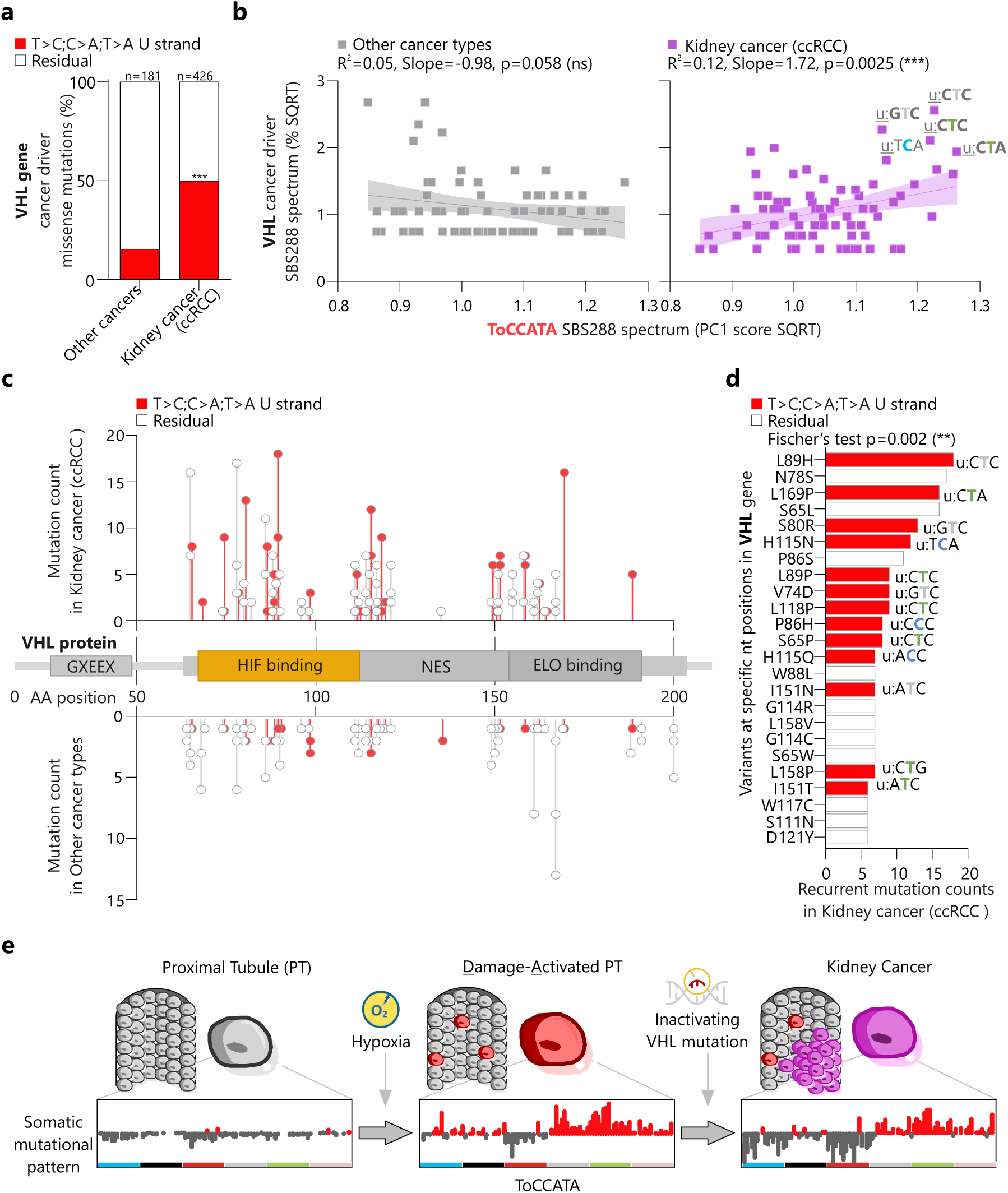
Functional contribution of ToCCATA to the generation of cancer driver mutations in ccRCC. **a**. Incidence of ToCCATA-related SBS types (U: T>C; C>A; T>A) on the SBS288 spectrum (shown in Suppl Fig 13a) of missense mutations labelled as “cancer driver” in the *VHL* gene in the AACR Project GENIE consortium database. Fischer’s exact test was used to compare the distribution of SBSs in ccRCC (exposed to the mutational process) vs other cancer types (not exposed to the mutational process) **** p<0.0001. **b.** Linear correlation of the SBS288 spectrum of *VHL* cancer driver variants with the SBS288 spectrum of ToCCATA (represented by the PC1 score of genic U: and T: SBS classes, shown in Fig 2c) in either ccRCC or other cancer types. Slope significantly different from zero was tested through simple linear regression. **c.** Schematics of the coding portion of the *VHL* gene and the nucleotides harboring cancer driver variants. The number of observed cases in the GENIE dataset is shown on the Y axis and the SBS types attributable to ToCCATA are represented in red. GXEEX: acidic repeat motif; HIF binding: Hypoxia inducible factor binding site; NES: nuclear export signal; ELO binding: Elongin C binding site. **d.** The top 25% most recurrent cancer-driver variants in *VHL* in kidney cancer (ccRCC). The specific SBS type and the relative aminoacid change are shown for mutations attributable to ToCCATA (highlighted in red). **e.** Graphical abstract depicting the transition from a normal PT to a damage-activated PT cell, marked by the ToCCATA signature, and the subsequent step towards kidney cancer (ccRCC) initiation. ** p<0.01; *** p<0.005, by Fischer’s exact test. ***p< 0.005; non-zero slope verified by simple linear regression.

We investigated which positions in the VHL sequence were more frequently mutated (Fig 5c). Specific nucleotide positions were recurrently harboring the same SBS (Fig 5c-d, Suppl Fig 13b). The frequently occurring variants (up to 18 recurrent cases in the dataset) were enriched for ToCCATA-related SBS types in ccRCC (Suppl Fig 13b). The variants with the highest numbers of co-occurrences in ccRCC (Fig 5d, the top quartile of Suppl Fig 13b) belonged to ToCCATA-related SBS classes in 57% of cases compared to 8% in other cancers (Fig 5d, Suppl Fig 13c), supporting a relevant impact of ToCCATA on the mutational landscape of ccRCC cancer driver variants.

Altogether, these findings indicate a functional role of the ToCCATA-related mutational process in kidney cancer initiation.

## Discussion

The first genetic event that drives carcinogenesis in ccRCC is the loss of function of the *VHL* gene ^22^, the negative regulator of adaptation to hypoxia ^41^. Oxygen levels are key to the kidney, which has one of the highest rates of oxygen consumption per unit tissue mass, second only to the heart ^42^. Elevated oxygen needs are mostly driven by the proximal tubule (PT), whose physiological activities are exceptionally energy-demanding ^38,43^. Here we show that both exposure to hypoxia and genetic loss of *VHL* impact the ability of PT cells to maintain genome stability. This role of the hypoxic pathway in genome maintenance is an important additional element to explain why adaptation to hypoxia is a core mechanism of tumor initiation in ccRCC.

Our high confidence, single genome, somatic mutation data of normal kidney cells detail a sequence of events that influence the transition from normal to cancer (Fig 5e). We discovered that a sub-population of PT cells that present transcriptional signatures of PT damage, depict a specific pattern of mutations (ToCCATA, Fig 2). The ToCCATA mutational signature is almost ubiquitously found in ccRCC genomes (Fig 2b), thus appearing as a necessary feature of ccRCC evolution. Nonetheless, the presence of the signature is not sufficient for tumor initiation. In fact, when growing cells directly from kidney biopsies, we found ToCCATA-bearing cells in the majority of healthy volunteers (5 in 6 when analyzing kidney biopsies from cancer-free individuals with normal kidney function ^28^). Nonetheless, only 1 in 38 males and 1 in 68 females develop ccRCC during their life ^44^. We consequently conclude that the ToCCATA-bearing cells are the population of kidney cells susceptible to evolve into ccRCC, but also that such transition requires additional steps.

We do not know the prevalence of *the DA-PT* cells in the kidney tissue, as our data are limited to a selected kidney-derived population that is able to expand *in vitro*. Our cell-culture based method dis-favors the growth of fully differentiated kidney cells, and instead enriches for tubular cells in a proliferative state. Single cell RNAseq studies of human kidneys have identified a *failed repair-PT* (also named *VCAM1-PT*) population that is activated by damage but fails to return to a fully differentiated state and remains in a chronically activated and proliferative state ^45^. This population has a frequency of around 4% of PT cells and increases with age ^12^. Most kidney-derived clones in our dataset show the transcriptional signature of failed-repair PT cells (Suppl Fig 1), suggesting that our culturing protocol enriches for this cell subset. However, among the 152 kidney clones that we tested by WGS, only 32% show the ToCCATA mutational signature, suggesting that only a fraction of *VCAM1-PT* cells is actually subjected to the mutational process inducing ToCCATA. In agreement, we have cases of ToCCATA-pos and ToCCATA-neg clones derived from the same individual and both showing the *VCAM1-PT* transcriptional profile.

ToCCATA is not just a marker of the population susceptible to become ccRCC. In fact, we provide evidence that ToCCATA directly favors tumor evolution, by increasing the chances of occurrence of cancer-driver, single nucleotide variants in the *VHL* gene (Fig 5). The ToCCATA signature specifically affects highly expressed genes (Fig 2) and this is an uncommon feature for a mutational process. In fact, transcribed DNA is subjected to more efficient repair and the predominant pattern is a reduction of mutation densities in transcribed regions ^36,46^. Accumulation of mutations in highly transcribed DNA in the *DA-PT* cells might be due to the formation of endogenous mutagens, for example, PT-specific by-products of cell metabolism ^6,8^. Moreover, we show that *in vitro* exposure to hypoxia is sufficient to induce some features of the ToCCATA signature. Hypoxia is a known challenge for genome integrity, altering the expression of multiple DNA repair pathway components ^47^, as well as the availability of metabolites, such as oxygen and alpha-ketoglutarate, cofactors of enzymes involved in genome maintenance and chromatin organization ^48–50^. Eviscerating the cellular mechanisms of mutations induced by oxygen perturbations in the kidney will require further work. Nonetheless, our finding that hypoxia drives kidney-specific mutagenesis may offer a mechanistic basis for the observation that metabolic and vascular comorbidities — obesity, type 2 diabetes, hypertension, CKD or the metabolic syndrome — are established risk factors for renal cancer with strong impact on incidence and mortality, since all these conditions converge on chronic or intermittent hypoxia of the proximal tubule ^1^.

An additional important finding of our study is that acute loss of *VHL* accelerates mutation accumulation (Fig 4). The activity of pVHL is recognized as a central node preventing kidney carcinogenesis ^41^ and biallelic loss of *VHL* has been recognized as the first and truncal event in ccRCC evolution ^21,22,51^. Nonetheless, loss of *VHL* alone is not advantageous for cell growth *in vivo*, in human PT ^52^, and in cell culture ^53,54^. Cultured PT cells lacking *VHL* undergo growth arrest, even in hypoxic environment ^54^. Therefore, *VHL*-loss likely confers alternative features that promote cancer. We provide *in vitro* evidence that the acute loss of *VHL* challenges genome integrity. Our data indicate that the role of *VHL* in genome maintenance is not restricted to protection from ToCCATA. For example, we observed that the loss of *VHL* sensitizes to the accumulation of oxygen-induced mutations during *in vitro* culture (Suppl Fig 12g), possibly suggesting a general hypersensitivity to DNA-damaging agents. A whole-genome Crispr screening has identified the VHL-HIF pathway as a node enabling correct repair of double-strand breaks ^55^. Moreover, a body of evidence has documented the presence of *VHL* in the nucleus and shown that *VHL* activity is essential for correct DNA repair and DNA synthesis under conditions of replicative stress ^56,57^. Interestingly, growth arrest in *VHL*-deficient cells could be rescued by concomitant loss of the *PBRM1* gene, a chromatin regulator, known subclonal driver in ccRCC ^56^. Therefore, our findings support the idea that *VHL*-loss directly promotes genome instability and that this pro-oncogenic event creates a platform that favors the occurrence of additional cancer driver mutations.

In summary, our high confidence detection of somatic mutations in normal cells from human kidneys provides unprecedented resolution of the transition from normal to cancer in this organ. We show that hypoxic episodes and loss of *VHL* converge into a vicious axis and promote genome instability, overall determining the early steps of carcinogenesis. Moreover, we propose hypoxic damage as a relevant physio-pathological factor stimulating tumorigenesis in the kidney PT.

## Methods

### Subject recruitment and specimen collection

Recruitment and collection of samples (blood, urine, tumor and normal kidney tissue biopsies) from either healthy volunteers (n=9, no history of kidney diseases and kidney cancer), VHL syndrome patients (n=9, with a genetic diagnosis and heterogeneous kidney cancer history, managed within the San Raffaele Hospital clinical program ^23^) or patients with sporadic kidney cancer and no genetic predisposition (n=1) was accomplished in agreement with the SoMuKT clinical study protocol, approved by the San Raffaele Hospital ethical committee (opinion 133/INT/2022) and amended on 25/08/2025. All enrolled patients provided written informed consent.

For each subject, we collected one blood sample and one to four urine samples. Reiterated urine collections from each individual spanned a period of 1-3 years. Biopsies of normal kidney parenchyma were collected from kidney nephrectomies occurring as part of oncological treatment. Tumor biopsies were obtained intraoperatively from 3 VHL-disease patients, undergoing either partial or radical nephrectomy. All samples were collected and sequenced within another study ongoing at the urological unit of San Raffaele Hospital (URBBAN). For two cases, WGS data of tumors, as well as other molecular and clinical information have been previously published ^58,59^. When WGS data from multiple biopsies from the same tumor were available, only one biopsy per tumor was included in this study, selecting the one with the highest tumor purity (as determined by WGS, see methods below). Samples with tumor purity <20% were discarded.

### Isolation and clonal expansion of kidney cells from human urines

One to four fresh urine samples (30-400 ml) were collected from each individual during regular urination or from the collection bag of a urinary catheter, and immediately processed. The protocol for cell isolation and culturing was adapted from ^60^. Briefly, urine samples were centrifuged and washed in PBS. After a second centrifugation, cells were resuspended in DMEM/F12 (GIBCO) containing 10% FBS and Renal Epithelial Cell Growth Medium (REGM) BulletKit (Lonza) and plated in gelatin-coated dishes (Merck). To obtain a clonal culture of single kidney cells, the first plating was done in a 24-well plate, which allowed for the growth of less than 1 colony per well in more than half of the urine samples. The 4^th^ day after plating, cells were shifted to REGM medium (Lonza) conditioned with REGM Bullet Kit (Lonza) including 0.5% FBS, and inspected weekly. Only distinct single colonies were considered for further culturing. When reaching 2-10 mm diameter, colonies were detached with trypsin, manually picked, and each colony was moved to a new gelatin-coated 6-well plate. Roughly 2/3 of the colonies expanded sufficiently to extract RNA, while 1/3 expanded enough for concomitant RNA and DNA extraction. Rare clones that expanded further were stored for additional experiments.

### Isolation and clonal expansion of kidney cells from human kidney biopsies

Three biopsies of normal kidney parenchyma were dissected by an expert pathologist, collected in PBS, and immediately processed for cell isolation and plating. Histological analyses confirmed the absence of cancer cells. Isolation of kidney epithelial cells was accomplished as in ^28^. Biopsy-derived colonies were expanded using the same protocol described for urine-derived cells, but medium was supplemented with 10% FBS.

### RNA extraction and qPCR

Total RNA was extracted using the Illustra RNAspin Mini kit (Cytiva), according to the manufactureŕs instructions. To minimize the batch effect of different RNA extractions and qPCR experiments, total RNA from one, normal kidney tissue, was repeatedly extracted in each experiment and used as a reference for relative quantification of gene expression. Each batch also included at least one sample of non-kidney cells, obtained from different preparations of human bone marrow-derived mesenchymal cells, subcutaneous fat progenitors, and skin fibroblasts, kept in culture for at least 3 passages. These samples were used as negative controls to quantify the baseline expression of kidney-specific genes in non-kidney cells. cDNA synthesis was performed using ImProm-II™ Reverse Transcriptase (Promega), and quantitative RT-PCR was performed using iTaq™ Universal SYBR® Green Supermix (BioRad). The quantification of target gene expression was normalized to the reference gene *H3*. To amplify each transcript, the following sets of primers were used:

**Table 1.**
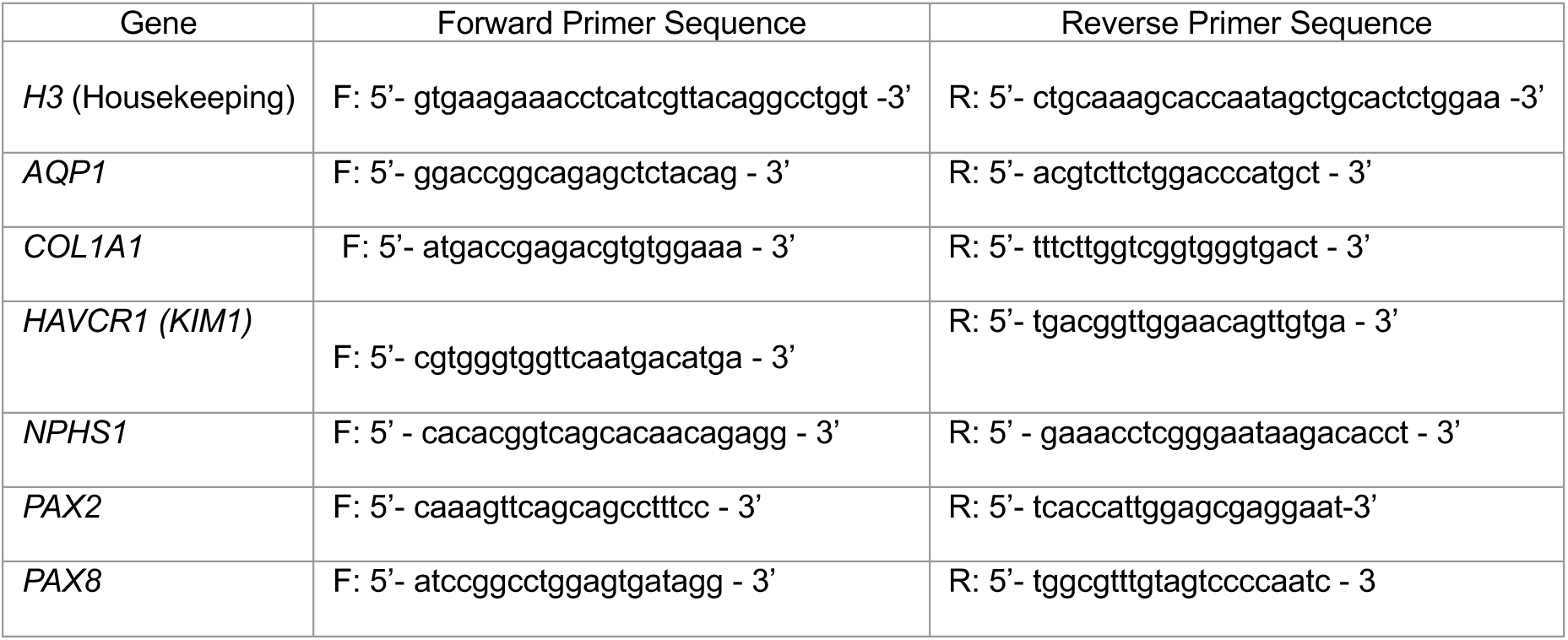
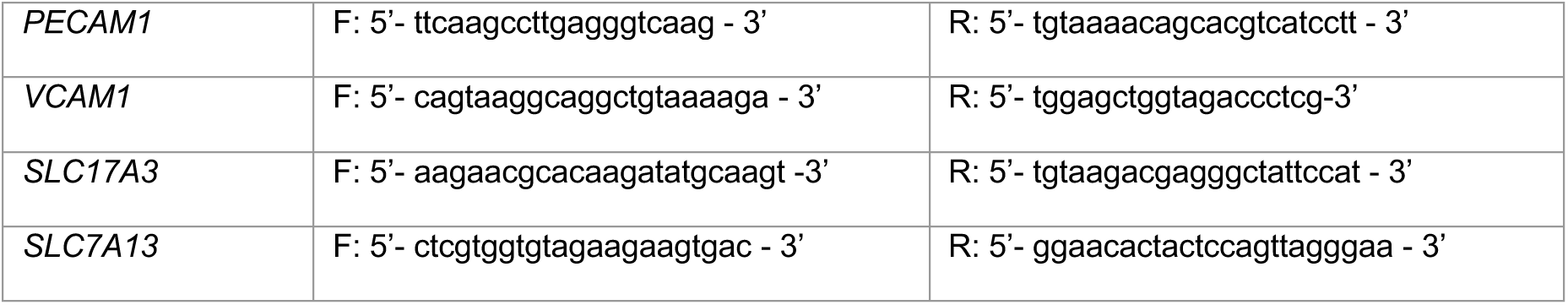
QPCR primers for gene expression analysis.

### RNA sequencing and QC

Two batches of RNA samples from kidney-derived clones (n=126 samples, minimum 200 ng total RNA/sample) were sequenced at Novogene Europe (Cambridge, UK). Briefly, RNA samples that passed QC control on sample quantitation, sample integrity (RIN) and sample purity (performed on Agilent 5400 fragment analyzer) were used for library preparation using the Novogene NGS RNA Library Prep Set (PT042). The quantified library was sequenced on the Illumina NovaSeq XPlus platform (PE150). Raw reads were filtered to remove adapter sequences, poly-N, and low-quality reads using Novogene in-house scripts. Quality was evaluated by Q20, Q30, and GC content metrics. Clean reads were aligned to the reference genome using HISAT2 (v2.0.5; ^61^), and read counts were assigned with featureCounts (v1.5.0-p3, ^62^). We obtained on average 20M reads assigned to genes and 14k expressed genes (expressed gene: 1 cpm in at least 2 samples) per sample. Plotting of the data by multidimensional reduction showed a sequencing batch effect, which was corrected using ComBat ^63^. Moreover, three outliers in the multidimensional reduction plot, as well as three samples that displayed abnormally high library size were excluded. The final dataset (n=120 samples) and their covariates are presented in Suppl Data 2. The UMAP plots of Suppl Fig 2 have been produced using the first 20 principal components.

The RNAseq data of HK2 clones (VHL-WT and VHL-MUT), used for expression-related mutation density analyses, are from ^39^.

### Transcriptional signature scores

Transcriptional signatures of 13 kidney cell types from human kidney biopsies were obtained from ^12^; while transcriptional signatures of 23 clusters of parenchymal kidney cells from human urines were obtained from ^17^. Briefly, specific markers from every cluster in either ^12^ or ^17^ were selected (p_val_adj < 0.05, log2FC >1 and pct.1 > 0.1) and then used to build signatures using ssGSEA to score them in 120 RNAseq samples from our dataset. The scores are presented in Supplementary Table 4. Suppl Fig 1b applies the same method, but using supra-cluster signatures, as presented in Fig S6 of the reference paper ^17^, as follows: Podocytes, clusters 1-2, (podocytes and PDC-injured); TEC_Differentiated, clusters 3-10 (proximal tubule, thin limb, thick ascending limb, distal convoluted tubule, connecting tubule, collecting duct principal cells, collecting duct intercalated cells); TEC_Inflamed, clusters 11-15; TEC_Damaged, clusters 16-18; TEC_Proliferative, cluster 19; TEC_Progenitor, cluster 20-23

### DNA extraction

Clone DNA was extracted from confluent wells of 6-multiwell plates using the Gentra Puregen Kit, Qiagen, following the manufacturer’s instructions. Blood DNA was extracted using the same kit from 3 ml of fresh or frozen total blood collected in EDTA, using the Gentra Puregen Blood Kit. DNA was extracted from tumor biopsies using Maxwell RSC 48/16 instruments (Promega, Madison, WI, USA) with Maxwell® RSC blood DNA kit, properly adapted for tissue sample processing. Quality of DNA was assessed with Genomic DNA screentape on Agilent Tapestation and only samples with DIN>7 were used for WGS.

### Whole genome sequencing

Library preparation and sequencing were carried out at HSR sequencing facility, using novaseq6000 with a 150×2 paired-end sequencing (Illumina S4 Reagent Kit v1.5, 300 cycles). NGS-library were prepared using the Illumina DNA PCR-Free Prep, Tagmentation kit and 100-1000 ng of DNA as starting material. Clone DNA was sequenced at a minimum average depth of 15x (range, average). Blood and Tumor DNA was sequenced at 60x. FASTQ files generation and secondary analysis were performed using Illumina DRAGEN (Dynamic Read Analysis for GENomics) Bio-IT platform available through the cloud Base Space Sequence Hub (BSSH, illumina). In particular the Dragen germline application was used for blood DNA analysis, the Dragen Somatic application was used for tumors and clones analysis.

### Somatic variant calling

Calling of somatic single nucleotide variants, small insertions/deletions (InDels) and Copy Number Alterations (CNAs) was performed for each clone/tumor genome using the Dragen somatic pipeline (v4.2.7) of Illumina BSSH, using the blood WGS germline data from the corresponding individual as a reference (tumor-normal analysis). The GRCh37/hg19 assembly version was used to align clone and relative blood data, while the GRCh38 version was used for tumors. Only VCF-PASS filtered variants were retained for further analysis. For clones (not applied to tumor samples), additional filters were applied. Variants with allele frequency <0.3 were removed. In addition, variants common to more than one clone from the same individual were considered missed germline and removed. A series of additional quality check steps were applied, to discard unsuitable clones: <65% of the genome covered at <10x (too low coverage); allele frequency distribution peak <0.35 (the population is not clonal); >20% of variants are shared with a different clone from the same individual (two clones are derived from a recent common ancestor). QC metrics of included and discared clones are shown in Suppl Data 3.

Somatic Copy Number Alterations (sCNAs) were obtained using the same somatic pipeline from Illumina DRAGEN somatic, and validated using CNVpytor as an additional calling pipeline. Germline pathogenic variants in established hereditary cancer predisposition genes (Suppl Table 1) were assessed through a tertiary analysis workflow using the eVai platform (enGenome). Within this gene set (*APC, ATM, BAP1, BMPR1A, BRCA1, BRCA2, BRIP1, CDH1, CDK4, CDKN1B, CDKN2A, CHEK2, EPCAM, FH, FLCN, GREM1, MEN1, MET, MLH1, MSH2, MSH3, MSH6, MUTYH, NBN, NF1, NF2, NTHL1, PALB2, PMS2, PMS2CL, POLD1, POLE, PTEN, RAD51C, RAD51D, RET, RHBDF2, SDHB, SDHC, SDHD, SMAD4, STK11, TP53, TSC1, TSC2, VHL*), variants were prioritized and interpreted by integrating population allele frequency, predicted molecular consequence, gene–disease validity, clinical evidence from curated databases (including ClinVar), functional data, and evidence from the scientific literature. For genes in which hereditary cancer predisposition is established in an autosomal recessive inheritance model, isolated heterozygous pathogenic or likely pathogenic variants were not considered reportable findings in the absence of a second pathogenic variant. Final variant classification was performed according to the ACMG/AMP guidelines.

### Generation of mutational matrices

Mutational matrices (spectra) for SBS, InDels and DBS were generated using SigProfilerMatrixGenerator with default options (v1.2.12) ^64^

### SBS signatures inference and attribution to COSMIC signatures

To maximize biological relevance and prevent spurious assignments of COSMIC signatures, a subset of genomes with known mutational exposures from published studies was co-analyzed within the dataset. The analysis was performed on the entire dataset displayed in Suppl data 4/Suppl Fig 5 (n=266 genomes). De novo mutational signatures were extracted from each mutational matrix using SigProfilerExtractor with nndsvd_min initialization (NMF_init = “nndsvd_min”) and default parameters (v1.1.9) ^33^. NMF was applied with factorizations between k = 1 and k = 15 signatures; each factorization was repeated 100 times ^33^. Subsequently, COSMIC attribution was performed using SigProfilerExtractor with default parameters (v1.1.9) ^33^. To confirm the results, the SBS signatures were extracted with a different method: MuSiCal ^65^. De novo signature discovery with default parameters was followed by decomposition into linear combinations of recognized COSMIC v3.4 single-base substitution (SBS) signatures. Decomposition was performed using MuSiCal’s multinomial likelihood-based sparse non-negative least squares (NNLS) model, while contribution coefficients were derived via maximum-likelihood refitting using a bidirectional likelihood model, applying a matching threshold of 0.0001 and a refitting cutoff of 0.001.

### PCA of SBS288-spectra and ToCCATA index

SBSs were classified into 384 classes using SigProfilerMatrixGenerator. This classification accounts for: the type of nucleotide substitution, its trinucleotide context, and its location in intergenic (N:) / genic DNA. The genic region is further stratified in transcribed (T:), untranscribed (U:) strand. Mutations in bidirectionally transcribed DNA (B:) are not shown and were not used for the analysis, resulting in a SBS288 matrix. The scaled SBS288 matrix (frequency distribution normalized on mutation counts) of 152 non-cancer kidney samples described in Fig 2b and c was used for PCA analysis. The PCA score relative to the PC1 was plotted in a heatmap and used to determine the profile of the ToCCATA signature (T<C; C>A, T>A on U; strand). The ToCCATA index, instead, was calculated for each sample as follows: (no. U:T<C + U:C>A + U:T>A) - (no. T:T<C + T:C>A + T:T>A)/ tot tSBSs. For the meta-analysis of published datasets, samples from the studies listed in Suppl Data 5 were used, after removing single genomes with <1000 tot SBSs, which gave unstable results. Samples from the same tissue, but from different studies were grouped as per Suppl Data 5.

### Clonal variants detection in tumor samples

A filter on allele frequency was applied on all somatic SBSs for each tumor and only variants with AF>0.25 were retained as clonal.

### Expression-related mutation densities

The effect of gene expression on SBS and indel density in cancer and normal kidney samples, as well as in *in vitro* mutations accumulated in HK2 clones was calculated as follows: genes were divided into 5 bins (quintiles, Q1-Q5), where Q5 and Q1 represented the 20% of genes with the highest and lowest expression in the kidney, respectively. For normal and cancer kidney samples, the ranking of genes based on gene expression was provided by a curated RNAseq database of normal kidney tissue (Human protein Atlas, http://proteinatlas.com). For the HK2 cells, the ranking was done based on RNAseq data generated in ^39^ and specifically refers to VHL-WT (n=4 replicates) or VHL-KO (n=4 replicates) samples in control oxygen. The mutation density in each quintile was calculated as the number of mutations divided by each region aggregate width measured in megabases. Values were normalized to the mutation density in the respective intergenic DNA and plotted for each cell type group reporting average and s.e.m.

The effect of gene expression on SBS288 classes, was obtained as follows: The number of mutations in Q1 was subtracted to the number observed in each of the other quintiles, for each SBS class. The resulting values were negative when mutation densities were higher in Q1 (absence of transcription) compared to transcribed DNA regions, overall highlighting the activity of transcription coupled repair (TCR). Conversely, positive values were obtained when transcribed regions had higher mutation densities, supporting the presence of transcription coupled damage (TCD). Mutation data from kidney and liver cancer were analyzed using the respective normal tissue gene expression data from HPA.

Similarity between mutational profiles was evaluated through cosine similarity computed in RStudio using the cosine function from the lsa package. The ToCCATA profiles defined in Fig 2e and 2g (defined as “ΔSBS/Mb”) were based on 192 SBS classes, as described. The cosine similarity scores were obtained by comparing: all 192 classes (genic); 96 SBS classes on the transcribed strand (T:); 96 SBS classes on the untranscribed strand (U:). For the ToCCATA mutation spectrum defined by PC1 scores in Fig 2c (defined as “PC1 ToCCATA”), the U: and T: components were tested separately, while the genic component includes both U: and T: (192 classes).

### Spearman correlation of ToCCATA index and gene expression

The lists (Suppl Data 6) of genes whose bulk RNA-seq expression robustly correlates with the ToCCATA mutational-signature score across kidney-derived clones, was obtained as follows and repeated in 3 datasets: all clones with available ToCCATA index and RNAseq (n=56, from 8 VHL+/+ and 3 VHL+/− individuals); individual VUD1 (n=11; *VHL+/−*); individual VUD12 (n=15; *VHL+/−*). For each dataset, genes were first filtered to those significant by univariate Spearman correlation (BH-FDR<0.05) and confirmed by multivariate PLS regression against ToCCATA (Variable Importance in Projection, VIP>1), producing the “robust gene” lists. For individual VUD12, no gene passed the BH FDR>0.05 filtering, therefore Suppl Data 6 reports the genes with p value <0.02 before correction. “Robust genes” were split by the sign of Spearman’s rho into a positively correlated set (expression rises with ToCCATA burden) and a negatively correlated set (expression falls with ToCCATA burden), and each set was tested separately for functional enrichment. Enrichment was computed with a hypergeometric (Fisher’s exact) over-representation test, BH-corrected within each library, against the GO_Biological_Process_2025 gene-set library. Terms in Fig 4b and e are reported with BH-adjusted p-values (pAdj).

### Distribution of cancer driver variants in *VHL*

Variants reported as “cancer driver” in the *VHL* gene by the the AACR Project GENIE Consortium ^40^ were sorted in two categories: “Kidney cancer (ccRCC)” and “Other cancers” (any other cancer type except renal and surrenal cancers). SBS288 spectra were obtained for each group of variants using SigProfilerMatrixGenerator, as described. Enrichment of ToCCATA-related variants (T>C; C>A, T>A on U; strand) was determined by Fisher exact test. Recurrent variants were sorted by counting the number of mutations (ToCCATA or not) causing the same amino acid variation. The most recurrent amino acid variations (ToCCATA or not) were compared between ccRCC and other cancers by Fisher exact test.

### Inducible knock-out of *VHL* in cell lines

Inducible KO of *VHL* was achieved in an immortalized, normal PT cell line (HK2 cells, ATCC # CRL-2190) *via* Crispr-Cas9 mediated knock-out of endogenous *VHL* accompanied by doxycycline (dox) inducible expression of a sgRNA-resistant, *VHL* transgene, as described in ^39^. Briefly, Cas9-expressing HK2 cells were first transduced with a dox-inducible, sgRNA-resistant *VHL cDNA* construct. Subsequently, knock out of the endogenous *VHL* was achieved by transduction of a *VHL*-targeting sgRNA and selection of 2 independent clones (MUT10 and MUT35), while a VHL wild-type clone (WT8) was obtained from cells transduced with a non-targeting control construct. After the necessary cell expansion passages, dox withdrawal stopped the expression of the transgene and allowed for complete and acute *VHL* deprivation exclusively in MUT10 and MUT35 clones *(VHL*-KO). Normal activity of the VHL protein under dox treatment, as well as the efficient depletion of the protein at 6 days after dox removal were tested by western blot. After assessing that WT8 cells with and without dox had comparable responses to hypoxia (both HIF1a nuclear levels, and SBS/indel counts), results from these two conditions were combined and presented as *VHL*-WT (n=6 subclones in control oxygen, including n=3 dox on and n=3 dox off; n=6 subclones in hypoxia, including n=3 dox on and n=3 dox off). Similarly results from two different VHL-KO clones (MUT10 and MUT35) were combined and constitute the *VHL*-KO group (n=4 subclones in control oxygen, including n=2 MUT10 and n=2 MUT35; n=4 subclones in hypoxia, including n=2 MUT10 and n=2 MUT35).

### *In vitro* experiments

Engineered HK2 cells were expanded in atmospheric oxygen (21-22% O2) unless otherwise stated. For hypoxia stimulation, cells were counted, seeded at low confluency in 6-well microtiters (2.5*10^4^ cells/well) and moved to a dedicated hypoxic workstation (Baker-Ruskinn SCI-tive Hypoxia Workstation), while an identical control plate was kept in the regular incubator. The hypoxia treatment consisted of 72h in 0.1% O2. The correct activation of the hypoxic pathway in low oxygen was tested by IF and WB, on material harvested before removing the cells from low oxygen. Upon treatment, the cells were moved to a bigger plate and expanded for 7 days. Ethylnitrosurea (ENU, Sigma N3385, 400 µM, compound dissolved in DMSO) was administered for 24h to VHL-WT HK2 (with dox), while cells were kept in regular atmospheric oxygen. No evident cytotoxic effect of ENU administration was noticed.

### Detection of *in vitro* mutations

To assess *in vitro* mutations in HK2 cells, treated cells were subcloned by limited dilution in a 96-well plate. 3-4 clones per condition were expanded in order to collect enough biological material to perform DNA extraction (30 days) and WGS, as described for primary cell clones. The somatic variant calling pipeline of the Illumina Dragen was run on each clone, using WGS data of the respective paternal cells (plate used for subcloning) as “germline”. This removed the variants that occurred in the HK2 cell line before Crispr-Cas9 editing and cloning. The allele frequency distribution peak of clones ranged between 0.32 and 0.34, instead of the regular 0.5 detected in primary cell clones. This reproducible peak is consistent with a state of aneuploidy of the immortalized HK2 cells. To remove subclonal variants, we filtered PASS variants with AF >0.225.

### Immunofluorescence

Primary kidney cells derived from urines and HK2 cells were fixed in PFA 4%, 0.1% Triton X-100 for 5 min and then blocked in 1% BSA for 1 hour. Permeabilized cells were incubated with indicated primary antibodies for 1 hour and fluorescent secondary antibodies for 1 hour. Cells were stained with DAPI and examined with Leica TSC-II SP8 confocal microscope. HIF1A (ab51608) was revealed using a fluorescent secondary antibody AlexaFluor488 (ab150077). DAPI staining was used to reveal the nuclei and measure nuclear translocation. Quantification of nuclear/cytoplasmic HIFa fluorescent intensity was achieved through a custom pipeline built in CellProfiler. A minimum of 20 cells per condition was counted.

### Western blot

Western blot analysis was performed on total cell lysate obtained through direct protein extraction in Laemmli buffer. The following antibodies were used: total H2A (Cell Signaling Technology #7631), H2AX (pS139) (Cell Signaling Technology #2577), HIF1A (Abcam ab51608), HIF2A (Cell Signaling Technology D9E3), beta-Actin (Cell Signaling Technology #4967), and VHL (Cell Signaling Technology #68547).

### Statistical analyses

Unless otherwise indicated, the p-values were calculated using either 2-tailed distribution, 2-sample unequal variance Student’s *t*-tests (when comparing 2 groups), while one-way or two-way Anova with multiple comparison post-hoc test in all the other cases. Significance was defined as p < 0.05 (*p < 0.05, **p < 0.01, ***p < 0.005, ****p < 0.001). The results are presented as the mean ± standard error of the mean (SEM). Calculations were performed using GraphPad Prism software. The linear fits between mutation numbers and age were obtained using a linear mixed-effects model where the dependent variable is the number of mutations, the fixed effect is age, and the random effect is the individual. Bonferroni correction was used to adjust for multiple testing. This analysis was performed using the lme4 package in RStudio.

## Data availability

The WGS raw data (FASTq) have been deposited at the European Genome Phenome Archive with Temporary Study ID 1781 (data transfer in progress). The RNAseq counts and the lists of somatic SNVs and indels produced in this study will be made available without restriction upon paper publication. Lists of somatic variants produced in the following studies: ^28,34,46,66,67^, can be downloaded from https://www.somamutdb.org/SomaMutDB/. Lists of somatic variants from ^68^ (normal liver) and ^37^ (kidney and liver cancers from PCAWG) are available with the cited publications.

## Code availability

Codes will be made available without restriction upon paper publication.

## Acknowledgments

We thank Maria Eriksson and Hafdis T Helgadottir for the 2019 dataset; Luigi Anastasia and Marco Piccoli for granting access to their Baker-Ruskinn SCI-tive Hypoxia Workstation; San Raffaele CUSSB (Centro Universitario di Statistica per le Scienze Biomediche) for statistical support.

## Author information

IF conceived the study. IF and JPM designed the experiments and obtained fundings. JPM, GBP, SF, FC, IR, VP, NS, GD, IF performed the experimental work. JPM, GBP, FC, VP, JMGM performed bioinformatics analysis. AL, IR, UC collected clinical samples. AL, AS, IF obtained ethical authorizations and conducted the clinical study. JG and SV provided the engineered HK2 cell lines. JPM, GBP, AL, SF, VP, NS, DMP, SV, FS, PC, AB, IF analyzed and interpreted the data. IF and JPM wrote the first draft of the manuscript and all authors critically revised the manuscript. All authors read and approved the final manuscript.

## Supplementary tables

**Supplementary table 1.** Subjects’ characteristics.

| VHL+/+ individuals |  |  |  |  |  |  |  |  |  |
| --- | --- | --- | --- | --- | --- | --- | --- | --- | --- |
| Acronym | Gender | Age | VHL pathogenic mutation | pVHL alteration | Germline pathogenic variants in established hereditary cancer predisposition genes * | Smoking status | Kidney disease | Kidney cancer | Other diseases and treatments |
| HUD6 | male | 24 | no | no | no | smoker | no | no |  |
| HUD4 | female | 29 | no | no | no | smoker | no | no |  |
| HUD1 | male | 40 | no | no | no | no | no | no | celiachy (untreated) |
| HUD13 | female | 42 | no | no | no | smoker | no | no |  |
| HUD3 | male | 52 | no | no | no |  | no | no |  |
| HUD5 | female | 53 | no | no | no | smoker | no | no | celiachy (treated); diabetes type 1 (treated) |
| HUD11 | female | 58 | no | no | no | no | no | no | metabolic syndrome (treated) |
| HUD2 | male | 63 | no | no | no | no | no | no | metabolic syndrome (treated) |
| STUD2 | female | 65 | no | no | no | n.a. | no | yes | nephrectomy for kidney cancer |
| HUD12 | male | 70 | no | no | no | smoker | no | no | prostate cancer (surveillance) |

| VHL+/- individuals |  |  |  |  |  |  |  |  |  |
| --- | --- | --- | --- | --- | --- | --- | --- | --- | --- |
| Acronym | Gender | Age | VHL pathogenic mutation | pVHL alteration | Germline pathogenic variants in established hereditary cancer predisposition genes * | Smoking status | Kidney disease | Kidney cancer | Other diseases |
| VUD6 | male | 29 | Chr3<br>10191506<br>NM_000551<br>.3:c.499C>T | Arg167Trp | n.a. | no | no | no | no |
| VUD29 | female | 30 | Chr3<br>10191570<br>NM_000551<br>.4:c.763T>C | Leu188Pro | no | no | no | yes | no |
| VUD17 | female | 35 | Chr3<br>10191506<br>NM_000551<br>.3:c.499C>T | Arg167Trp | n.a. | no | no | no | PCOS, DM1 |
| VUD12 | male | 39 | Chr3<br>10183864<br>NM_000551<br>.4:c.333C>G;<br>rs5030820 | Ser111Arg | n.a. | smoker | no | yes | DM |
| VUD11 | female | 42 | Chr3<br>10183864<br>NM_000551<br>.4:c.333C>G;<br>rs5030820 | Ser111Arg | n.a. | smoker | no | no | no |
| VUD9 | male | 44 | NM_000551<br>:<br>c.(?_1)_(64<br>2_?)del | Del full protein, Chr3p is retained | n.a. | smoker | no | yes | IPA |
| VUD4 | female | 48 | Chr3<br>10183857<br>NM_000551<br>.3:c.326delT | Del truncated protein p.(Ile109Thrfs*50) | n.a. | no | no | no | thyroid hyperplasia (euthyroidism), emiparesis |
| VUD2 | male | 55 | Del Chr3p (partial) | Del full protein and neighbouring genes: BRK1 Del full protein; FANCD2, GHRL truncated protein | no | no | yes | yes | DM2, IPA, ipoacusia, IRC |
| VUD1 | male | 56 | Chr3<br>10183793<br>NM_000551<br>.3:c.262T>G | Trp88Gly | no | no | yes | yes | Previous bilateral radical nephrectomy - dialysis |

\*tested cancer predisposition genes: *APC, ATM, BAP1, BMPR1A, BRCA1, BRCA2, BRIP1, CDH1, CDK4, CDKN1B, CDKN2A, CHEK2, EPCAM, FH, FLCN, GREM1, MEN1, MET, MLH1, MSH2, MSH3, MSH6, MUTYH, NBN, NF1, NF2, NTHL1, PALB2, PMS2, PMS2CL, POLD1, POLE, PTEN, RAD51C, RAD51D, RET, RHBDF2, SDHB, SDHC, SDHD, SMAD4, STK11, TP53, TSC1, TSC2, VHL*

**Supplementary table 2.**
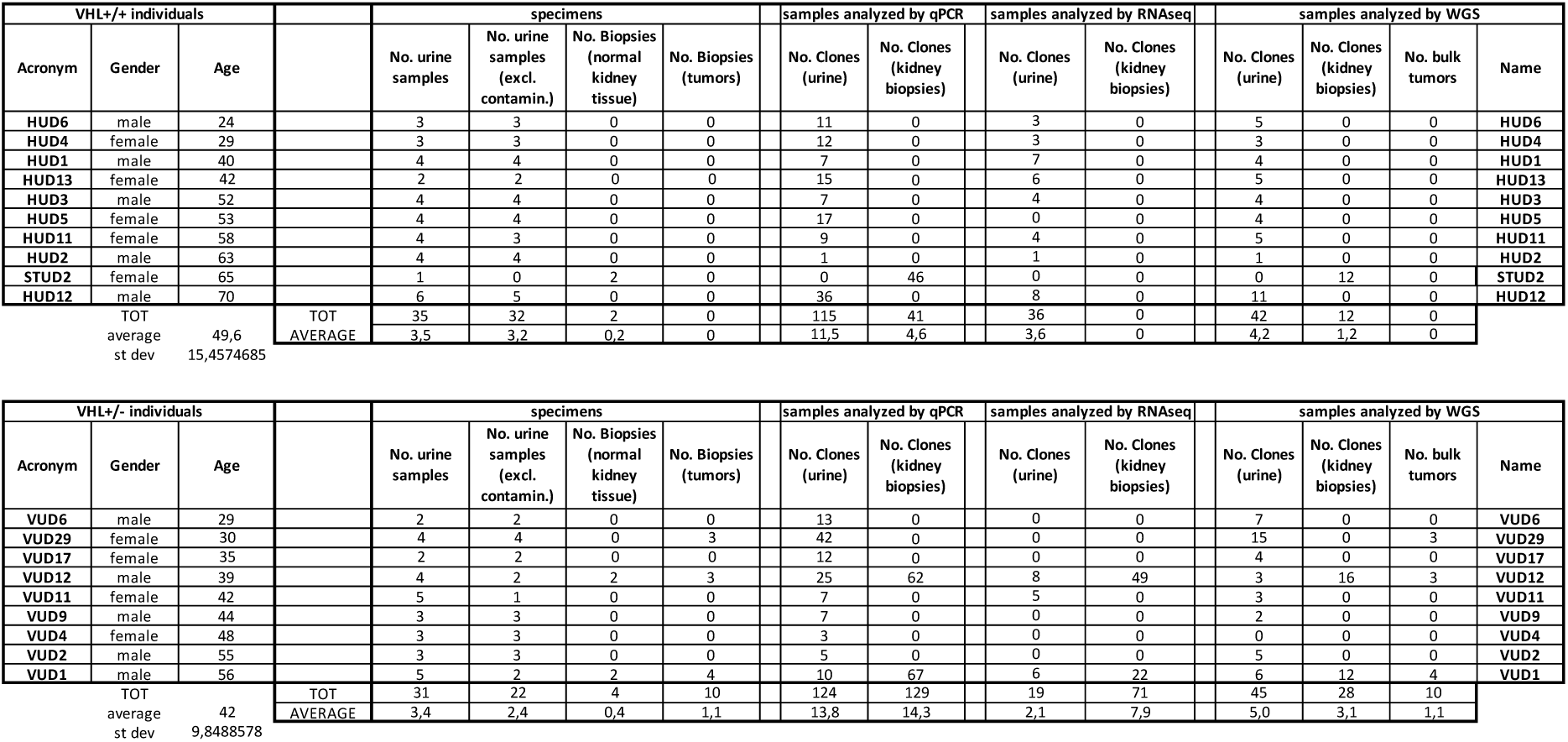
Analyzed samples per individual.

## Supplementary Data (separate file)

**Supplementary Data 1.** QPCR results per clone

**Supplementary Data 2.** RNAseq samples and signatures

**Supplementary Data 3.** WGS data per clone

**Supplementary Data 4.** COSMIC mutational signature analysis

**Supplementary Data 5.** Samples for ToCCATA index meta-analysis

**Supplementary Data 6.** RNAseq-ToCCATA Spearman correlation

## Supplementary figures

**Suppl Fig 1 (related to Fig 1):**
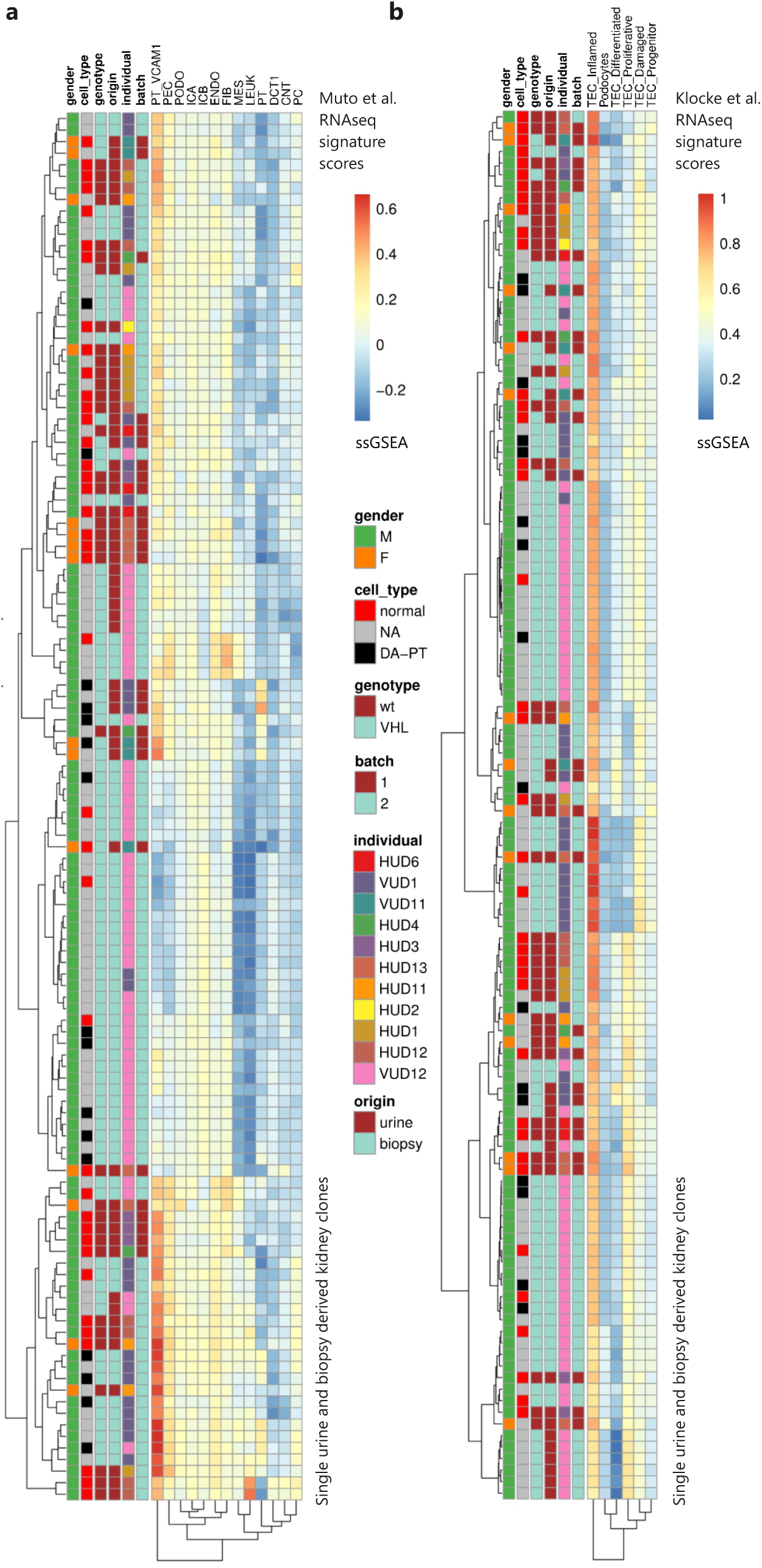
Kidney cell types transcriptional signature scores. **a.-b.** Heatmap depicting the ssGSEA scores of different kidney cell types transcriptional signatures for each kidney-derived clone (n=120). Transcriptional signatures were obtained from: **a.** ^12^ snRNAseq of kidney biopsies. PT, proximal tubule; PT_VCAM1, subpopulation of proximal tubule expressing *VCAM1*; PEC, parietal epithelial cells; TAL, thick ascending limb; DCT, distal convoluted tubule; CNT, connecting tubule; PC, principal cells, ICA, Type A intercalated cells; ICB, Type B intercalated cells; PODO, podocyte; ENDO, endothelial cells; MES, mesangial cells, FIB, fibroblasts; LEUK, leukocytes. **b.** ^17^, scRNAseq of urinary cells, supra-clusters as described in methods. TEC, tubule epithelial cells.

**Suppl Fig 2 (related to Fig 1):**
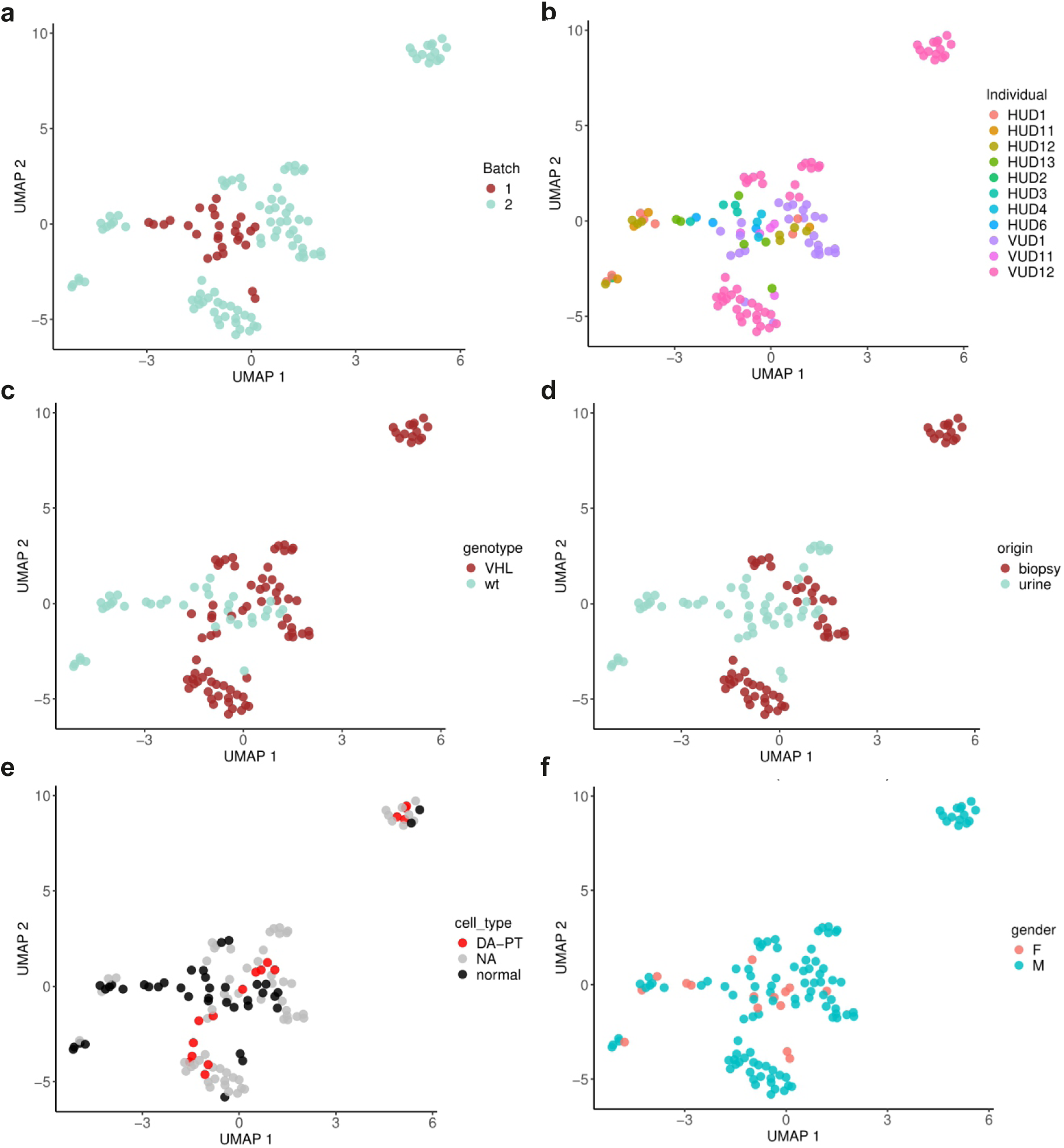
UMAP of bulk RNAseq from 120 kidney-derived clones. UMAPs depicting the nonlinear dimensionality reduction of kidney derived clones based on gene expression measured by RNA sequencing. Samples are highlighted in different colors, based on **(a)** RNA sequencing batch, **(b)** sample donor, **(c)** germline loss of function mutation in *VHL* (*VHL+/−)* or wild-type (*VHL+/+)*, **(d)** sample origin, **(e)** cell-type determined by the SBS288 principal component clustering of **Fig 1c**, **(f)** gender.

**Suppl Fig 3 (related to Fig 1):**
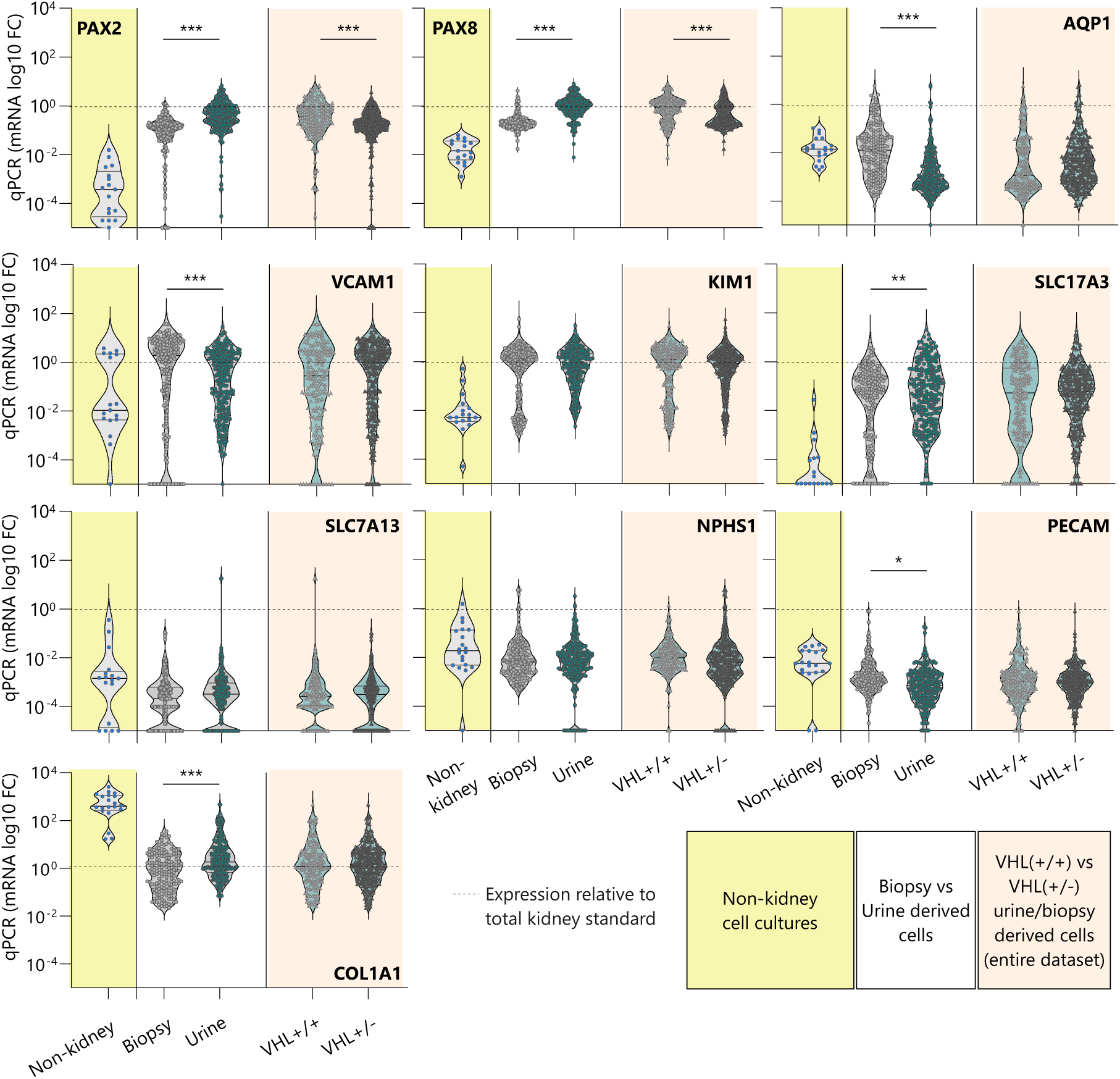
Quantitative PCR analysis of kidney marker gene expression in kidney-derived clones - Origin (kidney biopsy *vs* urine) and genotype (*VHL+/+ vs VHL+/−)* comparisons. **a**. Violin plots of the relative expression of kidney marker genes in different groups of normal kidney cell clones. Data from kidney-derived clones are compared with the expression levels quantified in cultured stromal cells derived from extrarenal tissues (bone-marrow derived mesenchymal cells, skin fibroblasts, fat progenitors). Expression levels were quantified by qPCR and reported as relative to a standard sample (normal kidney tissue biopsy). The panel shows the comparison between urine-(n=177) and kidney biopsy-(n=239) derived clones, as well as *VHL+/+* (n=162) vs *VHL+/−* (n=254) clones. * p<0.05; ** p<0.01; *** p<0.001, Student’s T test.

**Suppl Fig 4 (related to Fig 1):**
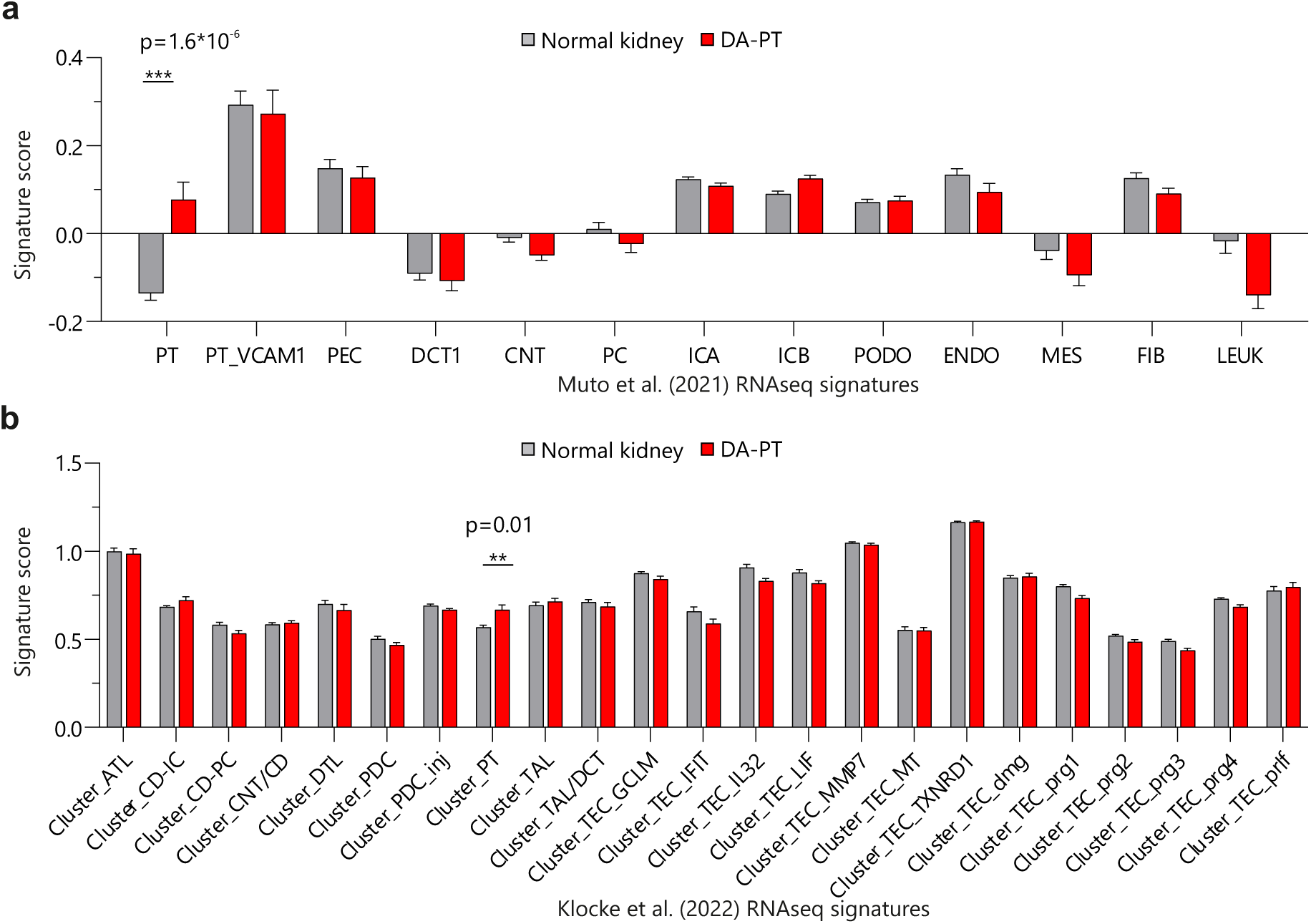
Kidney cell types transcriptional signature scores in DA-PT vs normal kidney cells. **a-b.** Average transcriptional signature score in DA-PT (n=17) vs normal kidney cells (n=40). Transcriptional signatures for the different kidney cell types were obtained either from ^12^ **(a.** snRNAseq of kidney biopsies) or from ^17^ **(b.** scRNAseq of urines**). a.** PT, proximal tubule; PT_VCAM1, subpopulation of proximal tubule expressing *VCAM1*; PEC, parietal epithelial cells; TAL, thick ascending limb; DCT, distal convoluted tubule; CNT, connecting tubule; PC, principal cells, ICA, Type A intercalated cells; ICB, Type B intercalated cells; ATL, thin limb, CD, collecting duct; IC, intercalated cell, PODO or PDC, podocyte; ENDO, endothelial cells; MES, mesangial cells, FIB, fibroblasts; LEUK, leukocytes. **b.** ATL, ascending thin limb; CD-IC, collecting duct intercalated cells; CD-PC, collecting duct principal cells; CNT/CD, connecting tubule / collecting duct; DTL, descending thin limb; PDC, podocytes; PDC_inj, injured podocytes; PT, proximal tubule; TAL, thick ascending limb; TAL/DCT, thick ascending limb / distal convoluted tubule; TEC_GCLM, tubular epithelial cells with glutamate-cysteine ligase modifier subunit expression; TEC_IFIT, tubular epithelial cells with interferon induced protein with tetratricopeptide repeats expression; TEC_IL32, tubular epithelial cells with interleukin 32 expression; TEC_LIF, tubular epithelial cells with leukemia inhibitory factor expression; TEC_MMP7, tubular epithelial cells with matrix metalloproteinase 7 expression; TEC_MT, tubular epithelial cells with metallothionein expression; TEC_TXNRD1, tubular epithelial cells with thioredoxin reductase 1 expression; TEC_dmg, damaged tubular epithelial cells; TEC_prg1, progenitor-like tubular epithelial cells subset 1; TEC_prg2, progenitor-like tubular epithelial cells subset 2; TEC_prg3, progenitor-like tubular epithelial cells subset 3; TEC_prg4, progenitor-like tubular epithelial cells subset 4; TEC_prlf, proliferating tubular epithelial cells. ** p<0.01; *** p<0.005; Student’s T test and Bonferroni correction for multiple testing.

**Suppl Fig 5 (related to Fig 1):**
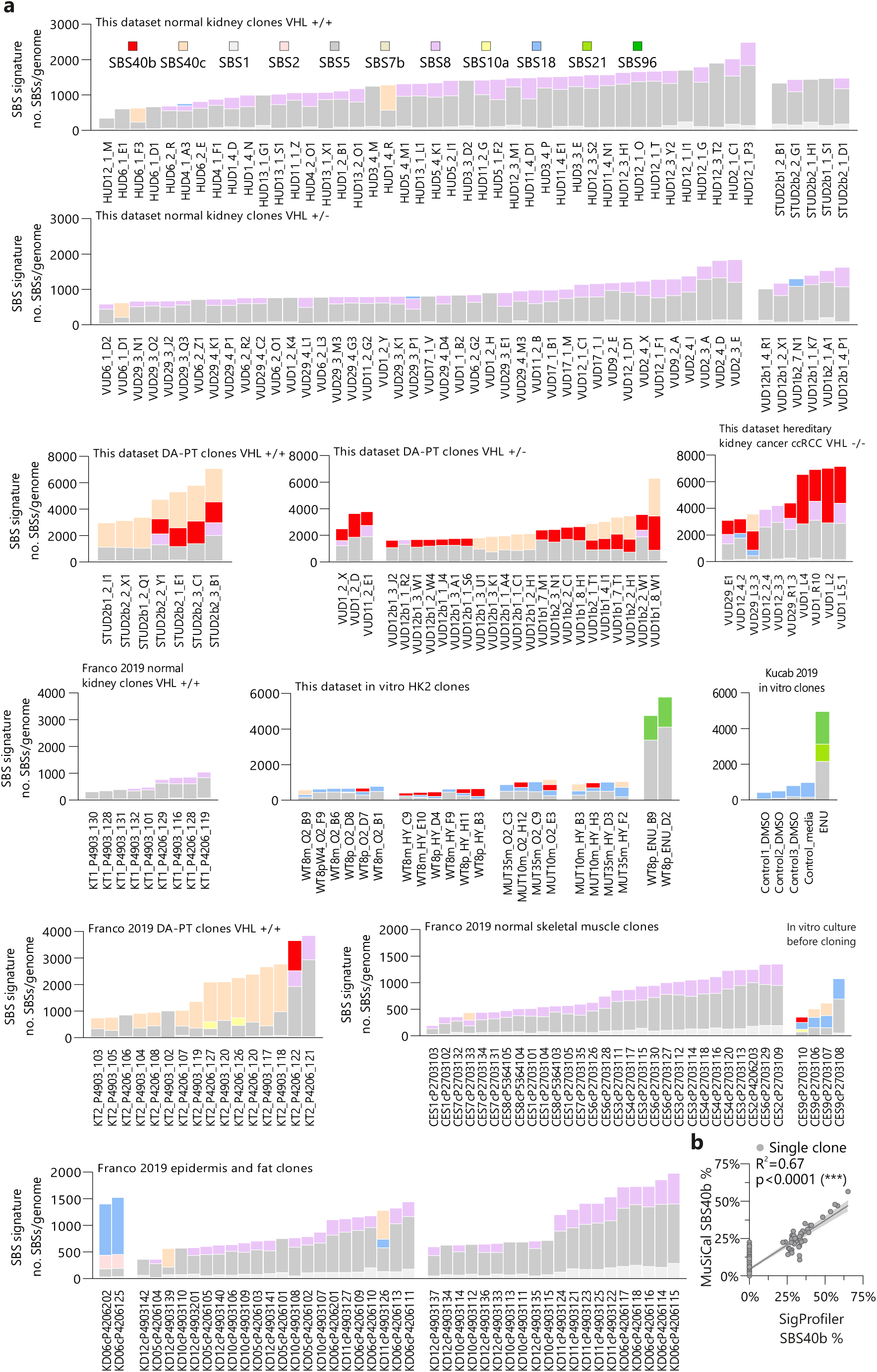
COSMIC SBS signature analysis in single genomes. **a.** Single genome results of COSMIC SBS signature analysis, including all samples shown in this study and a selection of published single genome data from ^28,34,35^. SBS signatures extraction and attribution to COSMIC signatures was done using SigProfiler. A list of samples included in the analysis and the numbers of SBSs attributed to each signature is reported in Suppl Data 4. **b.** Percentage of SBSs assigned to SBS40b in 266 samples shown in **a.,** using either SigProfiler or MuSiCal, and correlation of the two measurements in single clones ***p< 0.005; non-zero slope verified by simple

**Suppl Fig 6 (related to Fig 1):**
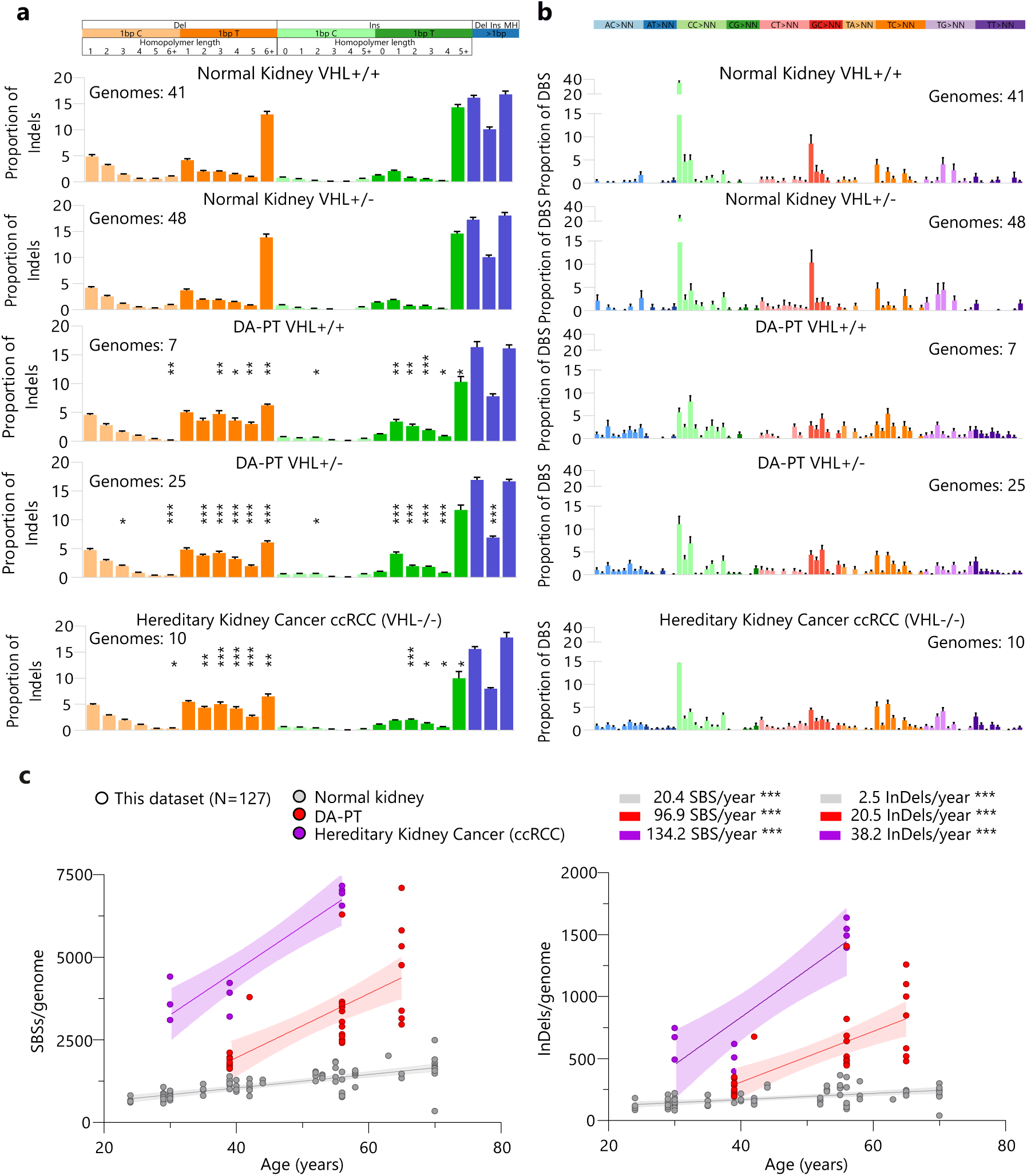
InDels and DBSs profiles in normal and cancer genomes from the kidney. **a.-b.** Somatic insertions and deletions (InDels) (**a**) and double base substitutions (DBSs) (**b**) profiles obtained using SigProfiler Matrix Generator. The average percentage and s.e.m. of InDels and DBSs attributed to each class is reported for indicated groups of normal and cancer genomes, together with sample numbers. **c.** Linear regressions of SBSs and InDels with age in VHL-syndrome kidney cancer (ccRCC), DA-PT and normal kidney cells. Intraindividual comparisons can be made for 3 VHL-disease patients, aged 29, 39 and 56, which have both normal and cancer genomes available. *p<0.05; ** p<0.01; *** p<0.005; by two-way ANOVA followed by Dunnett’s multiple comparisons test against the normal kidney VHL+/+ condition. ***p< 0.005; non-zero slope verified by simple linear regression.

**Suppl Fig 7 (related to Fig 1):**
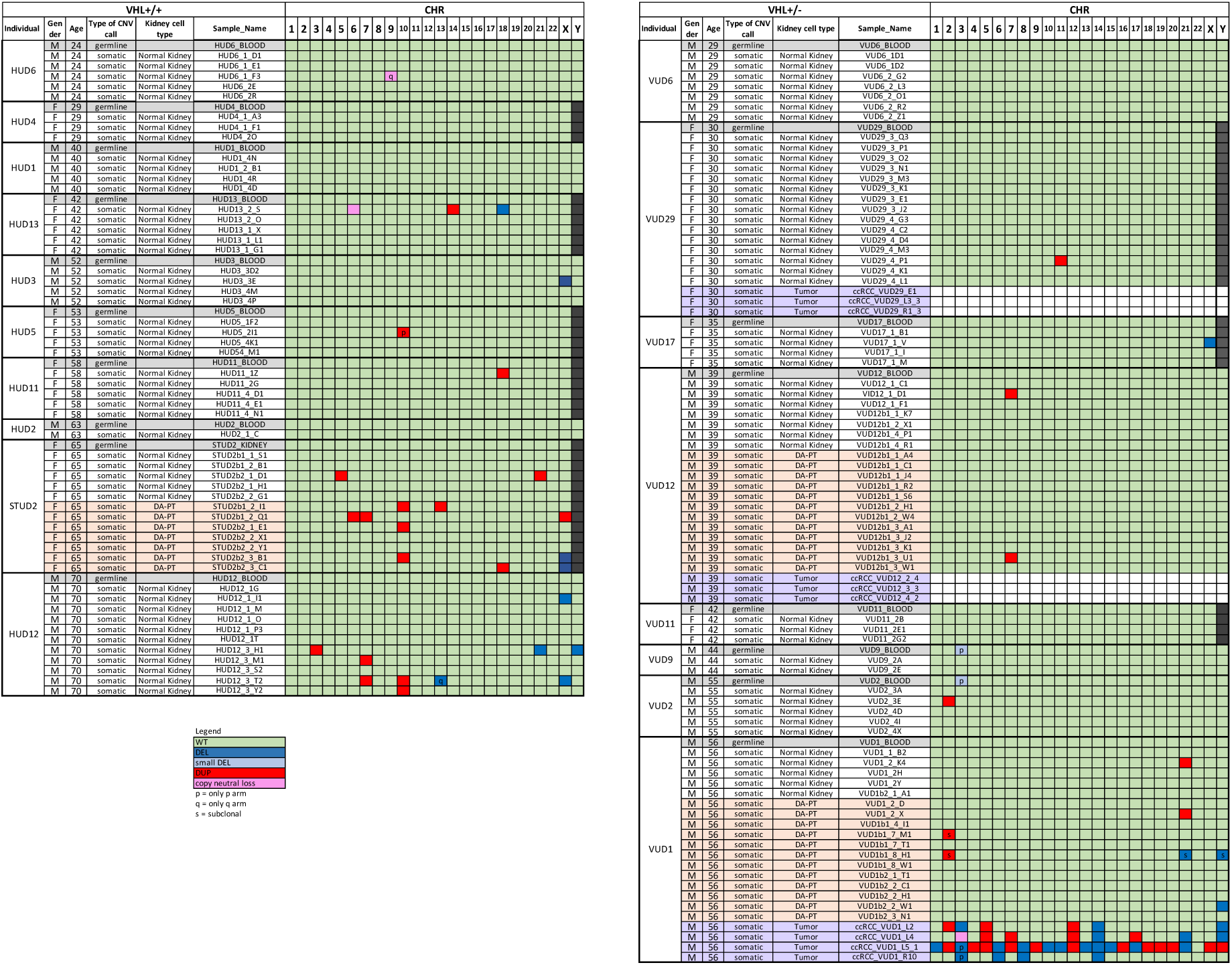
Major copy number variants in clones from VHL+/+ and VHL+/− individuals. Losses and gains of entire chromosomes or chromosome arms are reported for each individual germline (blood WGS analysis) and somatic genome (clone and tumor WGS analysis). For the somatic genomes, only the somatic events are reported (CNVs that are also found in the respective germline are not reported). Samples are grouped by VHL status (*VHL+/+* and *VHL+/−)* and individual. Samples are ordered by age and type of sample (*normal kidney*; *DA-PT,* highlighted in red; and tumor, highlighted in purple).

**Suppl Fig 8 (related to Fig 1):**
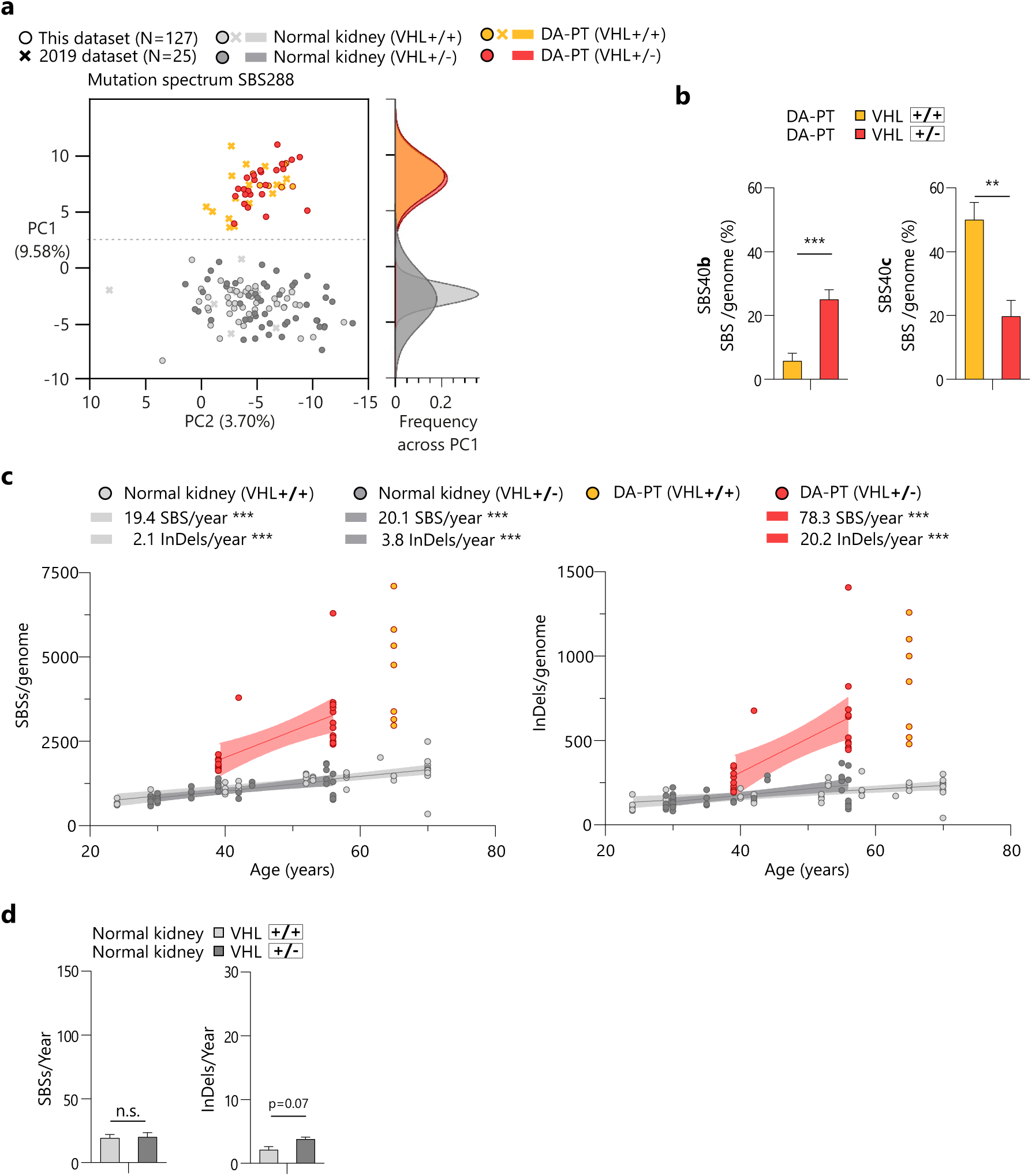
Comparison of somatic variants in genomes from VHL+/+ and VHL+/− individuals. **a.** PCA of SBS288 mutation spectra in 152 genomes from *in vitro* expanded *DA-PT* and *normal kidney cells*. The *VHL* status (*VHL+/+* and *VHL+/−)* of each genome is highlighted with different colours. **b.** Percentage of SBSs assigned to SBS40b and SBS40c in *VHL+/+* and *VHL+/− DA-PT* genomes (average with s.e.m.). **c.** Linear regressions and yearly increase of SBSs and indels with age in *DA-PT* and *normal kidney cells* from *VHL+/+* and *VHL+/−* individuals. **d.** Average SBSs and InDels per year with s.e.m. in normal kidney genomes from *VHL+/+* and *VHL+/−* donors. **p<0.01, ***p<0.005 by Student’s t-test. ***p< 0.005; non-zero slope verified by simple linear regression.

**Suppl Fig 9 (related to Fig 2):**
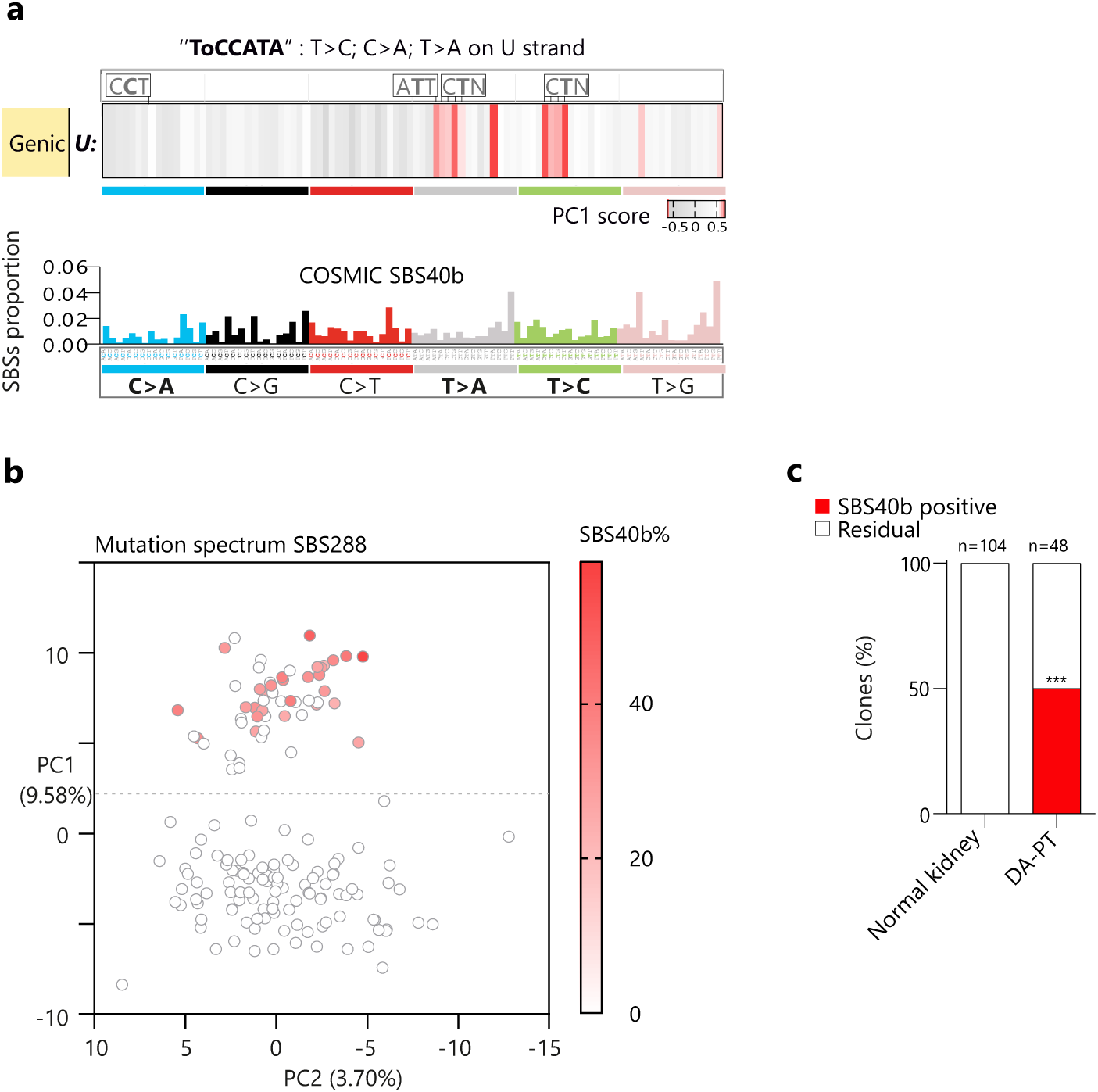
Comparison of SBS40b and ToCCATA. **a.** Spectrum of the COSMIC signature SBS40b (bottom) compared to the spectrum of ToCCATA (top), as obtained in Fig 2c. **b.** The percentage of somatic mutations attributed to SBS40b in our mutational signature analysis with SigProfiler (Supp Fig 5a) is represented by a red-color scale and overlayed to each dot of the PCA (Fig 1c) of the SBS288 spectra (representative of the ToCCATA signature) in single, non-cancer kidney genomes in this dataset. **c.** Fraction of SBS40b-positive genomes within the *DA-PT* versus *normal kidney cells* groups. *** p<0.005, by Fischer’s exact test.

**Suppl Fig 10 (related to Fig 2):**
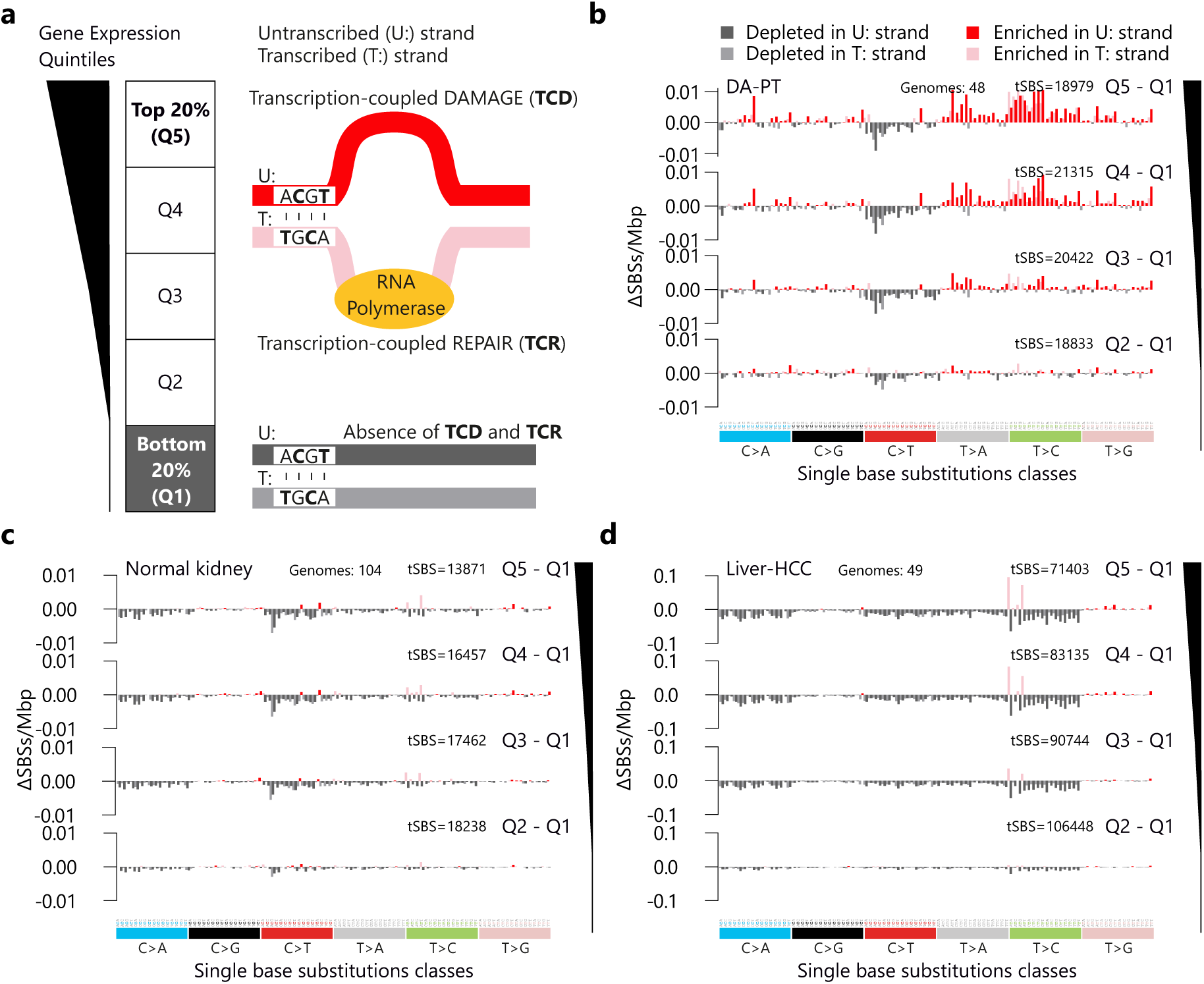
Transcription coupled mutation in kidney (non-cancer) and liver HCC. **a.** Schematic representation of the stratification of genic DNA into 5 quintiles, based on gene expression (Q1= undetectable expression; Q5=top expressed genes) and the differential activity of transcription-coupled DNA damage (TCD) and repair (TCR) in Q5 and Q1. **b.-d.** Histograms depicting the difference of mutational densities (mutations/Mbp) between transcriptionally active (Q5, Q4, Q3, Q2) and transcriptionally inactive (Q1) DNA for each SBS288 class. Negative values indicate a prevalence of TCR, while positive values correspond to TCD. Mutation data from DA-PT and normal kidney clones (n=48 and n=104 genomes, respectively, obtained from both urine/biopsy-derived cells from this study and biopsy-derived cells from the 2019 dataset ^28^) were stratified using consensus gene expression data of normal kidney tissue (Human Protein Atlas) (**b,c**). Difference of mutational density (mutations/Mbp) between the transcriptionally active and transcriptionally inactive DNA in liver cancer (hepatocellular carcinoma-HCC). n=49 genomes of HCC tumor biopsies ^37^. Genic DNA was stratified based on consensus gene expression data of normal liver tissue (Human Protein Atlas) (**d**).

**Suppl Fig. 11 (related to Fig 2):**
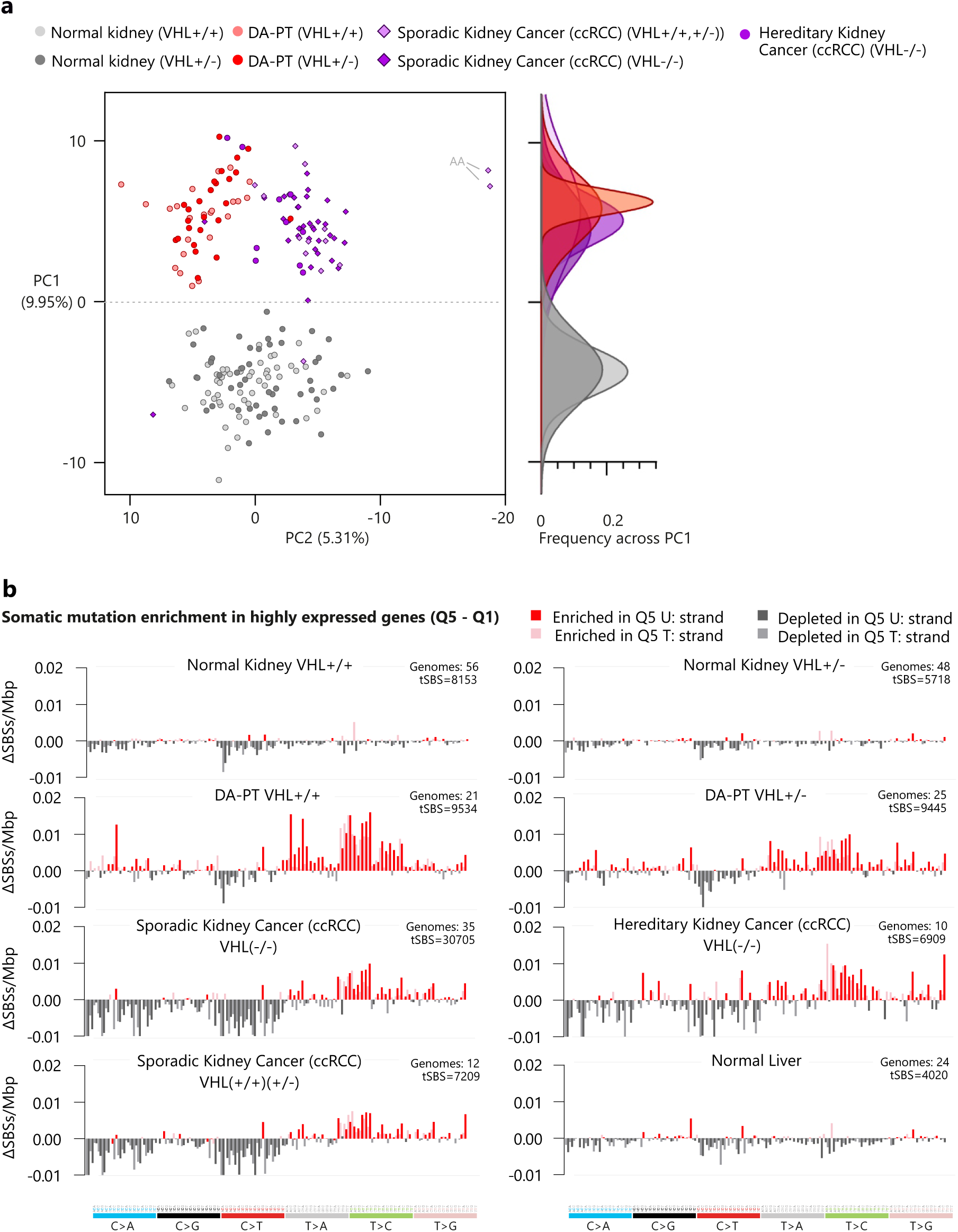
Effect of *VHL-*loss on mutation spectra and transcription-coupled mutation in normal and cancer kidney genomes. **a.** PCA of SBS288 mutation spectra in 201 genomes from sporadic and VHL-syndrome kidney cancers, and DA-PT and normal kidney clones (same as Fig 2f). Different groups are highlighted in different colours: *VHL* status (*VHL+/+* and *VHL+/−)* of each genome in *normal kidney* and *DA-PT cells*; sporadic (PCAWG) and inherited (VHL-syndrome) ccRCC. **b.** Spectrum of SBSs enriched in highly expressed (Q5) compared to non-expressed genic DNA (Q1) in aggregated somatic mutation data from cancer and normal genomes from kidney and liver. *VHL+/+* and *VHL+/−* genomes are analyzed separately. Sporadic and inherited ccRCC genomes are analyzed separately. For cleaner results, two sporadic ccRCCs showing aristolochic acid exposure have been removed. Normal liver and liver hepatocellular carcinoma (HCC) are shown.

**Suppl Fig. 12 (related to Fig 4):**
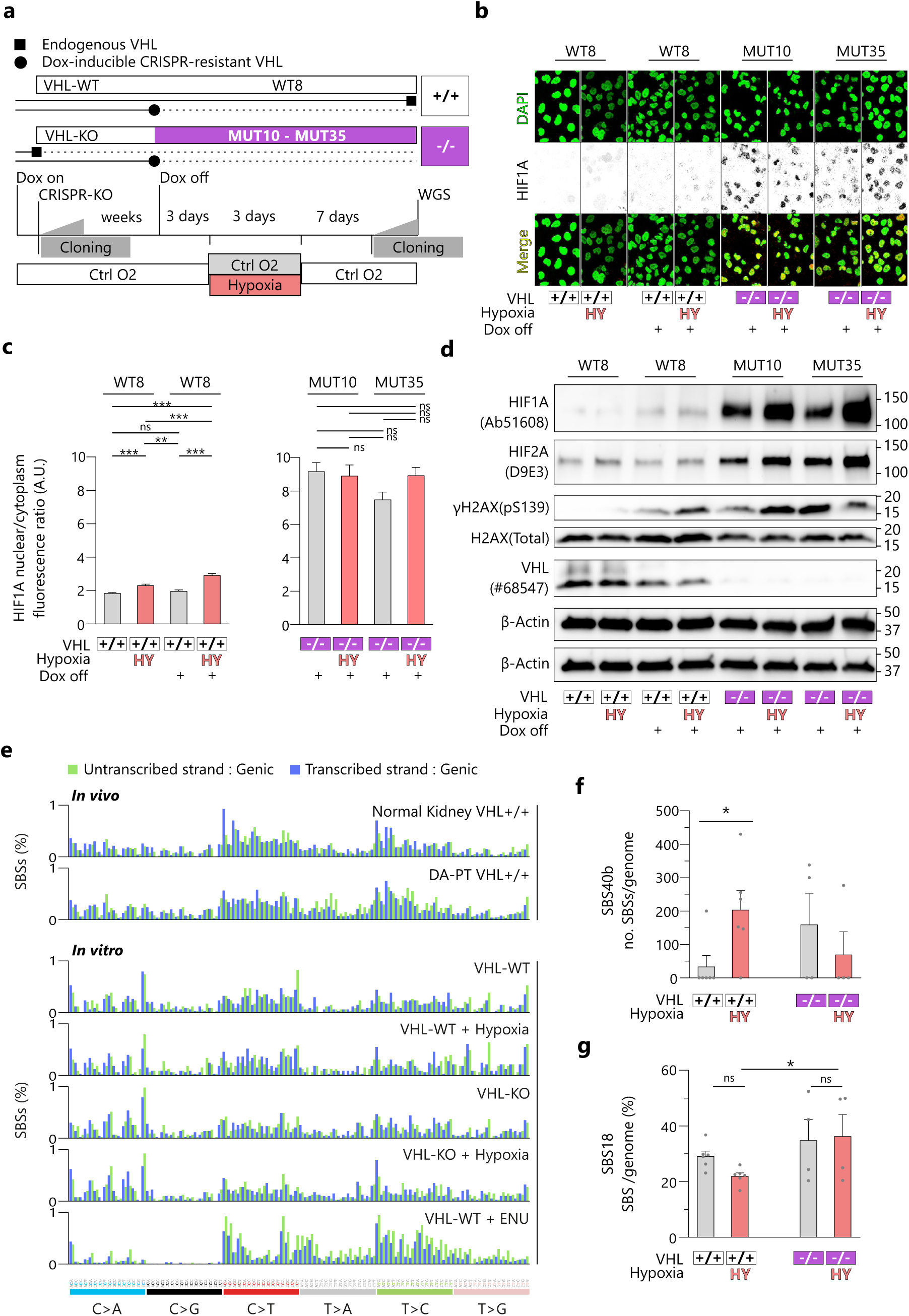
*In vitro mutagenesis* experiment in a normal PT cell line. **a.** Schematic representation of the strategy adopted to induce acute loss of *VHL* in the normal PT cell line HK2 and the experimental strategy for assessing *in vitro* mutation upon *VHL* loss and/or exposure to hypoxia (0.1% O_2_). **b.-d.** Representative images and quantification of IF (**b.-c.**) and WB (**d.**) analysis of the indicated proteins to measure the response to hypoxia in different lines of HK2 cells. WT8 cells (that underwent CRSPR-Cas9 with a non-targeting gRNA) were assessed in presence or absence of Dox. MUT10 and MUT35 cells (that underwent CRSPR-mediated KO of endogenous *VHL*) were assessed only after dox withdrawal and consequent stop of expression of the CRSPR-resistant *VHL* transgene. Cells were assessed immediately after exposure to hypoxia (0.1% oxygen). **e.** SBS288 genic profiles of *in vitro* mutations accumulated in HK2 cells cultured either in control oxygen (n=6 genomes) or exposed to putatively mutagenic stimuli: hypoxia (n=6), *VHL* deprivation (n=4), VHL deprivation + hypoxia (n=4), ENU 24h (n=2 genomes). The SBS288 profiles of *in vivo* mutations detected by cloning of primary kidney cells (either *normal kidney VHL+/+* n=9 or *DA-PT VHL+/+* n=14, from the 2019 dataset ^28^) have been added on top of the panel, to allow a comparison of *in vitro* and *in vivo* SBS profiles. **f.-g.** Counts of SBSs assigned to COSMIC SBS40b **(f.)** and percentage of SBSs assigned to SBS18 **(g.)** for each treatment group. *p<0.05; **p<0.01; ***p<0.005 by one-way ANOVA followed by Holm-Šídák’s multiple comparisons test, as appropriate.

**Suppl Fig. 13 (related to Fig 5):**
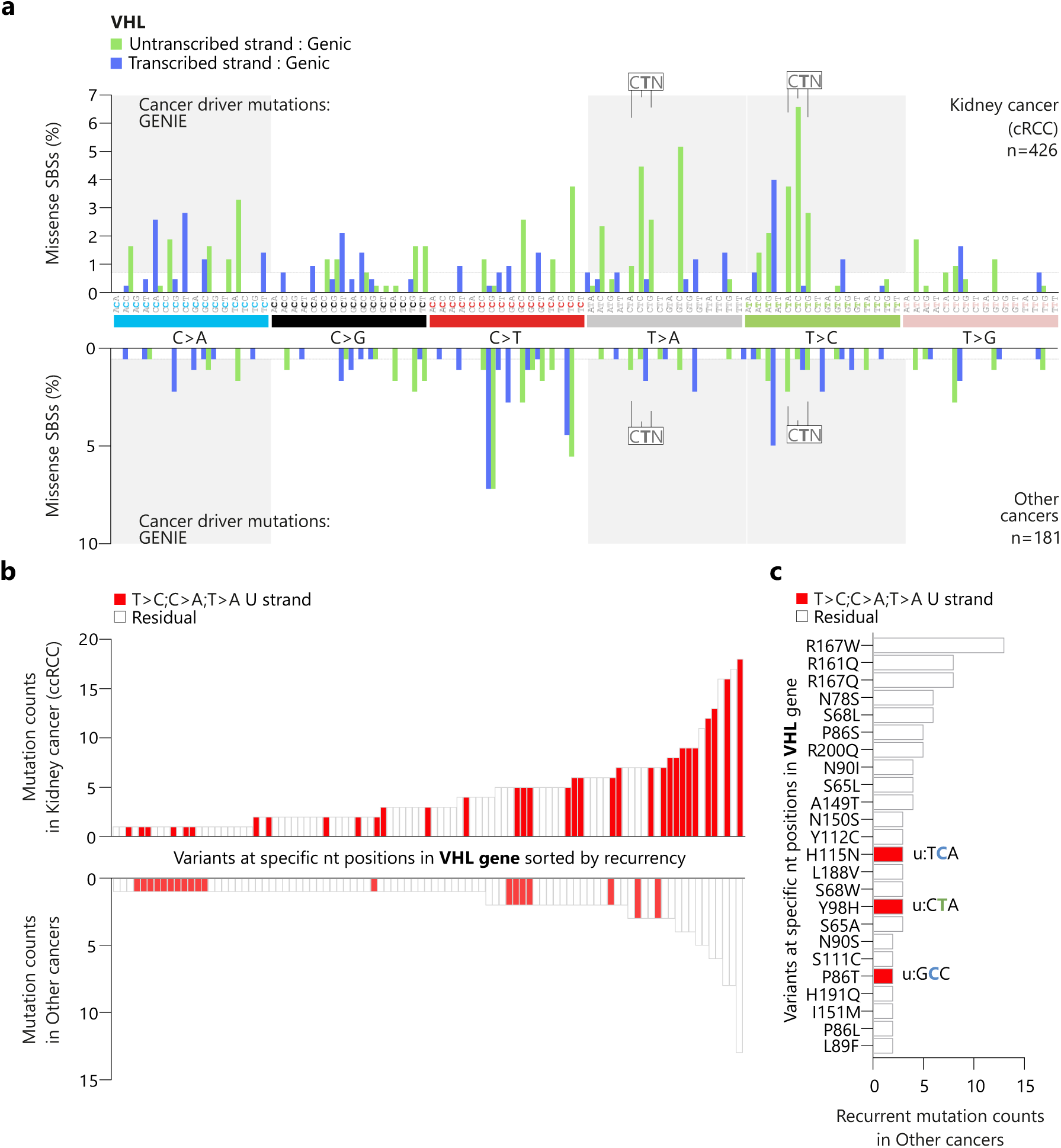
Functional role of ToCCATA in kidney cancer. **a**. Histogram depicting the SBS288 spectrum of missense mutations labelled as “cancer driver variants” in the *VHL* gene (AACR Project GENIE consortium database) found either in kidney cancer (ccRCC) (top, n=426 variants), or in any other cancers (bottom, n=181 variants). **b.** Ranking of *VHL* cancer driver variants, based on recurrency in the AACR Project GENIE consortium database for either ccRCC or other cancer types. **c.** The top 25% most recurrent cancer-driver variants in *VHL* in “other cancers” (any cancer excluding ccRCC). The specific SBS type and the relative aminoacid change are shown for ToCCATA mutations. Mutations attributable to ToCCATA are highlighted in red.

